# Projection Targeting with Phototagging to Study the Structure and Function of Retinal Ganglion Cells

**DOI:** 10.1101/2025.06.25.661576

**Authors:** Martin O. Bohlen, Andra M. Rudzite, Tierney B. Daw, Genevieve M. Kuczewski, Ergi Spiro, Cassie Hammond, Darienne R. Rogers, Alejandro Gallego-Ortega, Suva Roy, Kimberly Ritola, Marc A. Sommer, Greg D. Field

## Abstract

Visual information from the retina is sent to diverse targets throughout the brain by different retinal ganglion cells (RGCs). Much of our knowledge about the different RGC types and how they are routed to these brain targets is based on mice, largely due to the extensive library of genetically modified mouse lines. To alleviate the need for using genetically modified animal models for studying retinal projections, we developed a high-throughput approach called *projection targeting with phototagging* that combines retrograde viral labeling, optogenetic identification, functional characterization using multi-electrode arrays, and morphological analysis. This method enables the simultaneous investigation of projections, physiology, and structure-function relationships across dozens to hundreds of cells in a single experiment. We validated this method in rats by targeting RGCs projecting to the superior colliculus, revealing multiple functionally defined cell types that align with prior studies in mice. By integrating established techniques into a scalable workflow, this framework enables comparative investigations of visual circuits across species, expanding beyond genetically tractable models.

**Motivation:** Visual information from the retina is distributed to diverse targets throughout the brain. Much of our knowledge about how visual information is processed and routed to these brain targets is based on mice because of the large library of genetically modified mouse lines. For most other species, such libraries are not available. Therefore, we were motivated to develop an approach for characterizing diverse retinal projections into the brain that can be applied to other species. We aimed to achieve projection targeting with retrograde viral vectors, followed by identification of circuit-specific retinal ganglion cells (RGCs) with optogenetics, functional characterization with multi-electrode array (MEA) recordings, and morphological description with *in situ* and confocal microscopy. The resulting high-throughput approach permits functional and morphological investigation of dozens to hundreds of cells in individual experiments. By replacing the need for genetically modified animals with a suite of standard techniques, this approach should improve cross-species comparisons of the initial stages of visual processing.

**Summary:** Understanding the structure-function relationships across neurons is challenging, particularly when circuits are composed of dozens of distinct cell types. We refined an approach, called *projection targeting with phototagging,* that allows simultaneous elucidation of the projections, morphology, and visual response properties of diverse RGC types in the mammalian retina. The approach combines retrograde virally mediated phototagging of RGCs, microscopy, and large-scale MEA measurements. Importantly, the approach does not rely on transgenic animals and thus is generalizable across species. We validated this approach in rats by targeting retinal projections to the superior colliculus (SC). We showed that multiple RGC types project to the SC and that these results in rats align well with prior findings from transgenic mouse studies.

**Graphical abstract:** 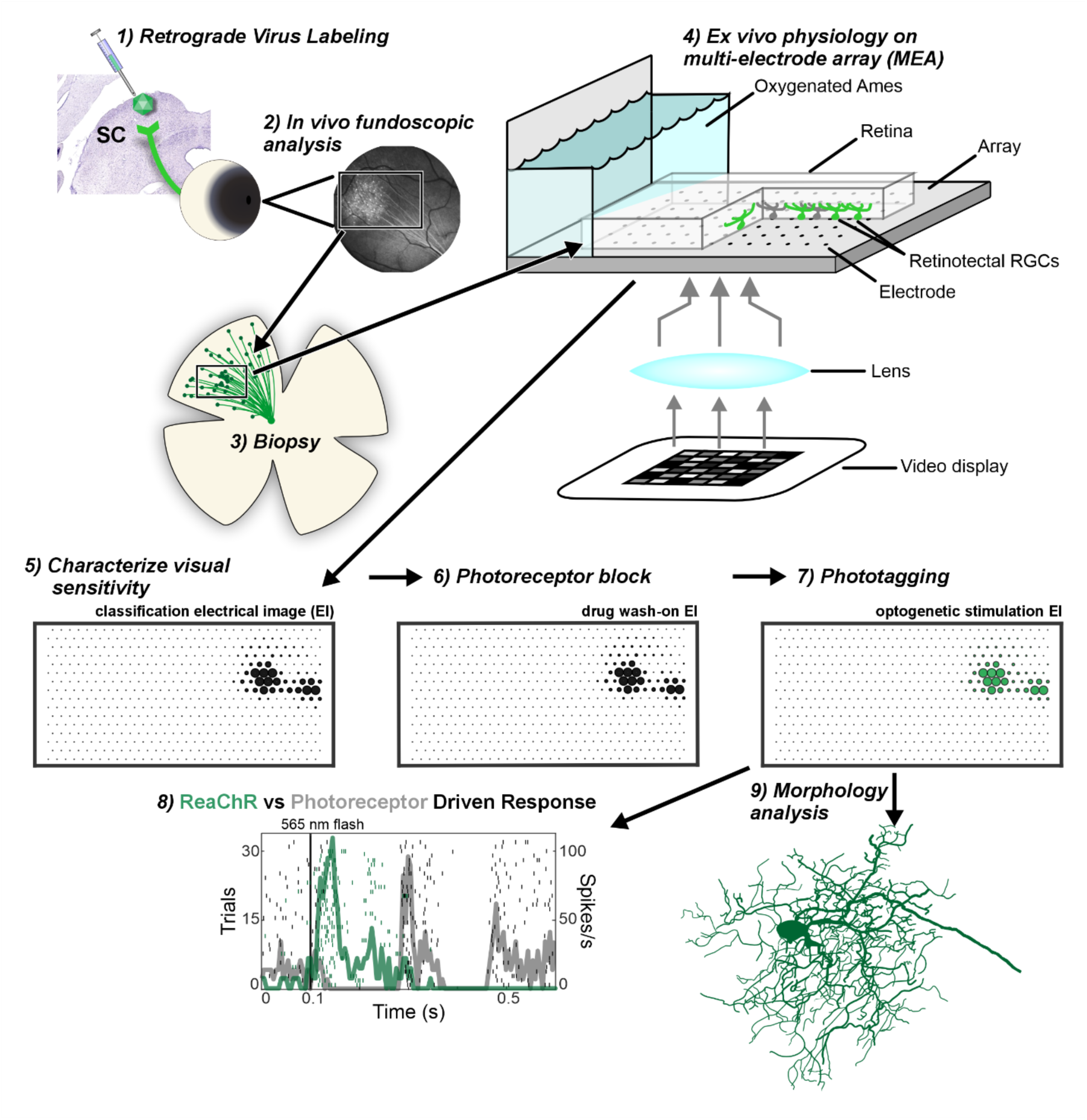

**Highlights:**

- The viral vector, AAV2retro, is effective for delivering opsins to retinal ganglion cells (RGCs) for phototagging and projection mapping.
- Phototagging allows matching RGC visual responses to morphology, while also identifying their central projections.
- The approach does not rely on genetically modified animals and thus is generalizable across species.

## INTRODUCTION

Understanding the relationship between structure and function is a central goal in neuroscience, as it illuminates how biological building blocks underpin information processing systems. The relevance of structure-function relationships is growing with the advent of high-throughput methods. For example, large-scale electrode recordings measure signals across thousands of neurons, and connectomics reconstructs neuronal morphologies and synapses within entire circuits. However, these functional and structural approaches are not frequently deployed within the same experiments, resulting in impressive morphological datasets that can be divorced from function or vice versa. Thus, high-throughput approaches that allow simultaneous assessment of cellular morphology, axonal projections, and neuronal signals are needed to generate a unified perspective on structure and function.

The retina is an ideal system for developing approaches that simultaneously elucidate structure, function, and projections across diverse cell types. Our knowledge of retinal projections has heavily relied on traditional tracers such as horseradish peroxidase and viral constructs like rabies (Itaya, 1980; Hofbauer and Drager, 1985; Ellis et al., 2016). More recently, the use of transgenic mouse lines has expanded the atlas of retinal ganglion cell (RGC) types and their specific projections (Huberman et al., 2008; Yonehara et al., 2008; Huberman et al., 2009; Kim et al., 2010; Bae et al., 2018; Goetz et al., 2022): current accounts indicate that the mouse retina contains ∼45 RGC types (Bae et al., 2018; Goetz et al., 2022) that send axons to ∼50 distinct brain areas (Martersteck et al., 2017). By contrast, studies in primate, cat, and rabbit retinae suggest fewer than 20 RGC types that send their axons to only ∼15-20 brain areas (Sanderson and Pearson, 1977; Klooster et al., 1983; Zhang and Hoffmann, 1993; Rockhill et al., 2002; Dacey et al., 2003; Yamada et al., 2005; Sivyer et al., 2011; Ellis et al., 2016; Peng et al., 2019; Kling et al., 2024). Are these species differences genuine, or do they reflect technological throughput differences between mice and other mammals? Answering this question will ultimately require methods that do not rely on genetically modified animals and that can be deployed across species.

To address this need, we refined a technique termed projection targeting with phototagging (also referred to as optotagging) to study retinal projections and RGC diversity. We applied this technique to the rat visual system, specifically to RGCs that project to the superior colliculus (SC). We injected the SC with an adeno-associated virus (AAV) that infects axon terminals at the injection site and is retrogradely transported to the cell body (AAV2retro). This virus drove the expression of two genes. One gene encoded a fluorescent protein that served as an anatomical marker for identifying transduced retinotectal RGCs. This provided a comprehensive intracellular fill, enabling high-resolution morphological reconstruction of individual RGCs. The other gene encoded the red-shifted channelrhodopsin (ReaChR)(Lin et al., 2013), which allowed us to match optogenetically driven responses to photoreceptor-mediated responses in RGCs. We confirmed the expression of virally mediated genes using *in vivo* fluorescence fundoscopy, then measured the spiking responses of RGCs from *ex vivo* retina using a large-scale multi-electrode array (MEA). Cumulatively, this approach allowed for accurate matching of the morphological profiles of individual RGCs and the visual signals they transmit to the SC without the need for genetically modified animals. Similar to results from transgenic mice, we found that phototagging in rats revealed a broad diversity of RGC types projecting to the SC, including cells with classical center-surround receptive fields and cells with orientation- and direction-selective preferences (Yonehara et al., 2008; Huberman et al., 2009; Ellis et al., 2016).

## RESULTS

### Viral targeting of SC projecting RGCs

The initial step towards identifying RGCs that project to the SC involved selectively targeting these cells for transgene expression. To achieve this, we employed the AAV2retro capsid (Tervo et al., 2016) and the CMV early enhancer/chicken β actin (CAG) promoter (Okabe et al., 1997; Alexopoulou et al., 2008) to drive expression of a red-shifted channelrhodopsin, ReaChR (Lin et al., 2013), and a fluorophore, mCitrine (Griesbeck et al., 2001). The virus was injected stereotactically into the superficial (Zo, SuG, Op), intermediate (InG, InWh), and/or deep (DpG, DpWh) tectal layers at the central third of the SC along the medial-lateral and rostral-caudal axes (**Fig. 1**).

**Figure 1:**
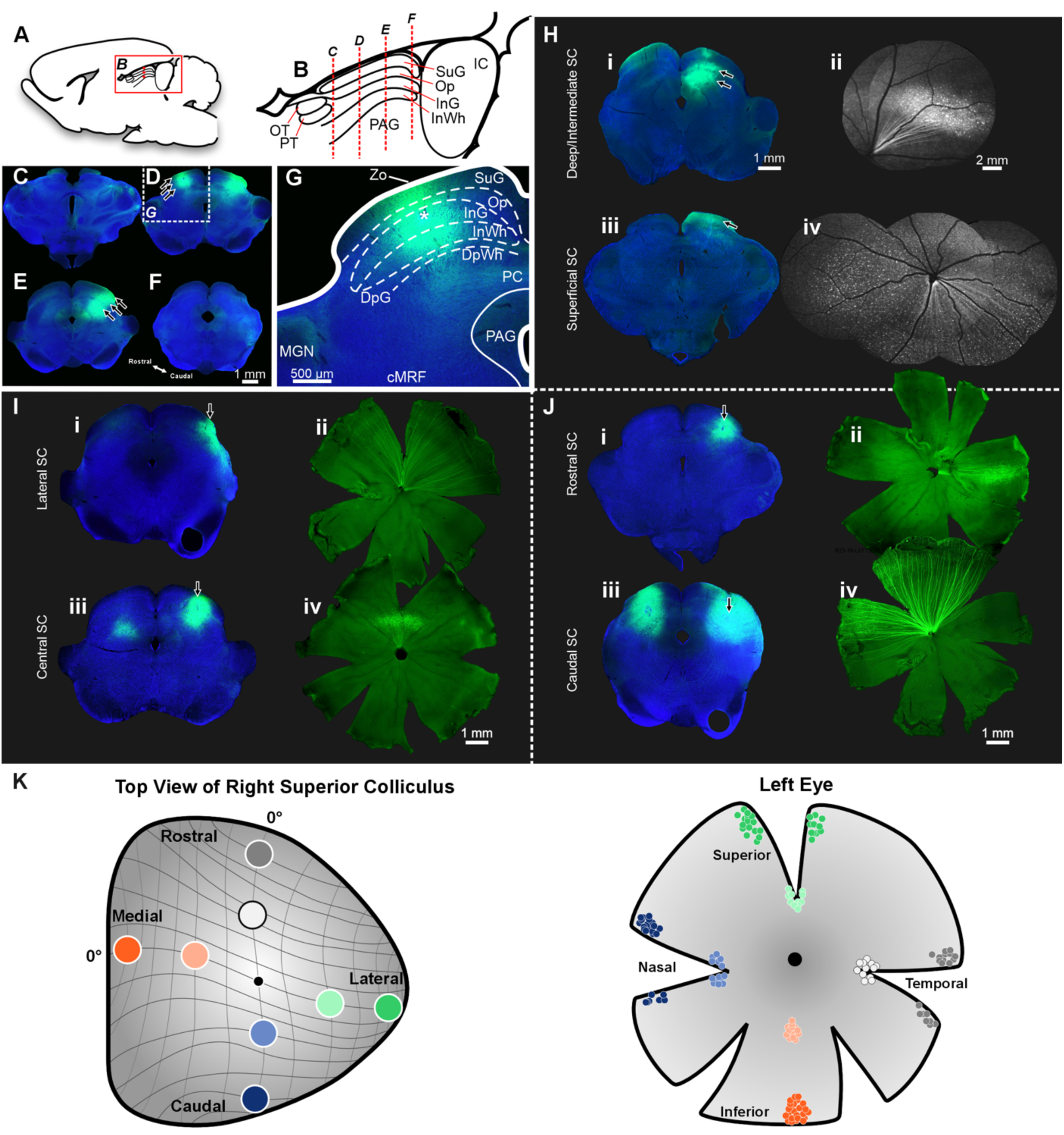
Targeting RGCs using retrograde virus. **(A)** Midline parasagittal drawing of the rat brain. Box illustrates the location shown in B. **(B)** Parasagittal schematic illustrating the laminar structure of the SC. Dotted red lines indicate the approximate locations of coronal sections shown in C-F photomicrographs. **(C-F)** Representative rostral **(C)** to caudal **(F)** digital photomicrographs of coronal sections through the midbrain for a rat that received bilateral AAV2retro-CAG-ReaChR-mCitrine injections at three depths. Arrows indicate locations of putative injection sites based on the presence of GFP-amplified mCitrine expression. Box in D indicates the location shown in G. **(G)** High magnification view of viral injections in the left SC with fluorescent labeling distributed across the intermediate and superficial layers. The * denotes a putative needle track. **(H)** Example cases illustrate different injection locations in the rat SC and the resultant RGC labeling. An injection into the intermediate and deep lamina of the tectum **(Hi)** resulted in focal RGC labeling **(Hii)**. An injection largely confined to the superficial tectal lamina **(Hiii)** resulted in diffuse RGC labeling **(Hiv)**. **(I)** Examples demonstrate retinal labeling arising from injections along the medial-lateral axis of the SC. An injection in lateral SC **(Ii)** resulted in peripheral superior RGC labeling **(Iii)**. In contrast, a more centrally placed injection (along the medial-lateral axis) into the tectum **(Iiii)** resulted in superior labeling more proximal to the optic nerve head **(Iiv)**. **(J)** Examples illustrate retinal labeling following injections along the rostral-caudal axis of the tectum. Injections into the rostral SC **(Ji)** resulted in temporal retinal labeling **(Jii)**, while caudal SC injections **(Jiii)** generated nasal retinal labeling **(Jiv)**. **(K)** The schematic depicts the right hemisphere SC (left panel) with representative injection sites (colored circles) and the contralateral retina (right panel) with resultant topographically organized labeled RGCs.

This approach builds on prior research demonstrating that AAV2retro can deliver transgenes to neuronal populations having projections to the tectum (Cushnie et al., 2020; Nanjappa et al., 2022; Daw et al., 2023; Zheng et al., 2024). In a representative case following bilateral injections, diffusion of the virus in the left tectum was confined to the intermediate and superficial layers (**Fig. 1D, G**), while in the right tectum the injection site had viral diffusion that extended into the deeper layers of the SC (**Fig. 1E**). Prominent labeling was present bilaterally in the superficial optic layers, where RGC axons heavily terminate (Hofbauer and Drager, 1985; May, 2006; Ellis et al., 2016) (**Fig. 1D, E, G**).

We then compared the viral expression pattern in the retina with the locations of injections in the SC. Injections focused on the intermediate and deep lamina of the tectum (**Fig. 1Hi**) resulted in punctate RGC labeling (**Fig. 1Hii**), whereas injections into the superficial lamina (**Fig. 1Hiii**) resulted in diffuse RGC labeling (**Fig. 1Hiv**). Along the medial-lateral axis, injections into the lateral aspect of the SC (**Fig. 1Ii**) resulted in far peripheral superior RGC labeling (**Fig. 1Iii**), while injections placed more centrally along the medial-lateral axis (**Fig. 1Iiii**) resulted in RGC labeling that was superior and proximal to the optic nerve head (**Fig. 1Iiv**). For these experiments, we avoided injections into the medial SC because this would label RGCs in the inferior retina (Drager and Hubel, 1976). We instead aimed for the superior retina because it contains cones expressing mostly M-opsin, which is ideal for eliciting physiological responses based on the emission spectra of our visual display (Galindo-Romero et al., 2022). Finally, rostral SC injections (**Fig. 1Ji**) resulted in temporal retinal labeling (**Fig. 1Jii**), while caudal SC injections (**Fig. 1Jiii**) generated nasal retinal labeling (**Fig. 1Jiv**). The pattern of RGC transduction following deposition of retrograde virus in different parts of the SC is summarized in Figure 1K.

### In vivo fundus imaging of RGC transduction

Some variability with the location and depth of the SC injections was present across animals as illustrated in the previous section (**Fig. 1**). Consequently, this led to differences in the precise location of labeled RGCs across retinas. Since the area of the MEA (∼2 mm^2^) is a small fraction of the retina’s total area, we needed a method to visualize the location of transduced cells prior to placing retinal segments on the MEA for *ex vivo* electrophysiology.

To identify the locations of labeled RGCs, we used *in vivo* fluorescence fundus imaging (Nanjappa et al., 2022), which provided several advantages. First, it allowed for a minimally invasive examination of transgene expression in the retina prior to sacrificing the animal, which was crucial for identifying and placing photosensitive and healthy retinal segments containing transduced RGCs on the MEA. In a few early experiments, fluorescent RGCs were instead imaged after enucleation, which resulted in substantial photobleaching and thus reduction in the sensitivity of physiological recordings. Fundoscopy performed several days before euthanasia ensured photoreceptors had recovered from any potential photobleaching during the fundus imaging and provided an accurate map of retinal locations with labeled RGCs (**Fig. 2, 3**). Fundus imaging also permitted multiple imaging sessions of the same retinas after injection, allowing measurement of the expression timeline (**Fig. 2**)(Nanjappa et al., 2022; Zheng et al., 2024). Between 2-3 weeks post-injection, the expression pattern in the retina stabilized (**Fig. 2E-L**) and remained constant for several months (**Fig. 2M-O**).

**Figure 2.**
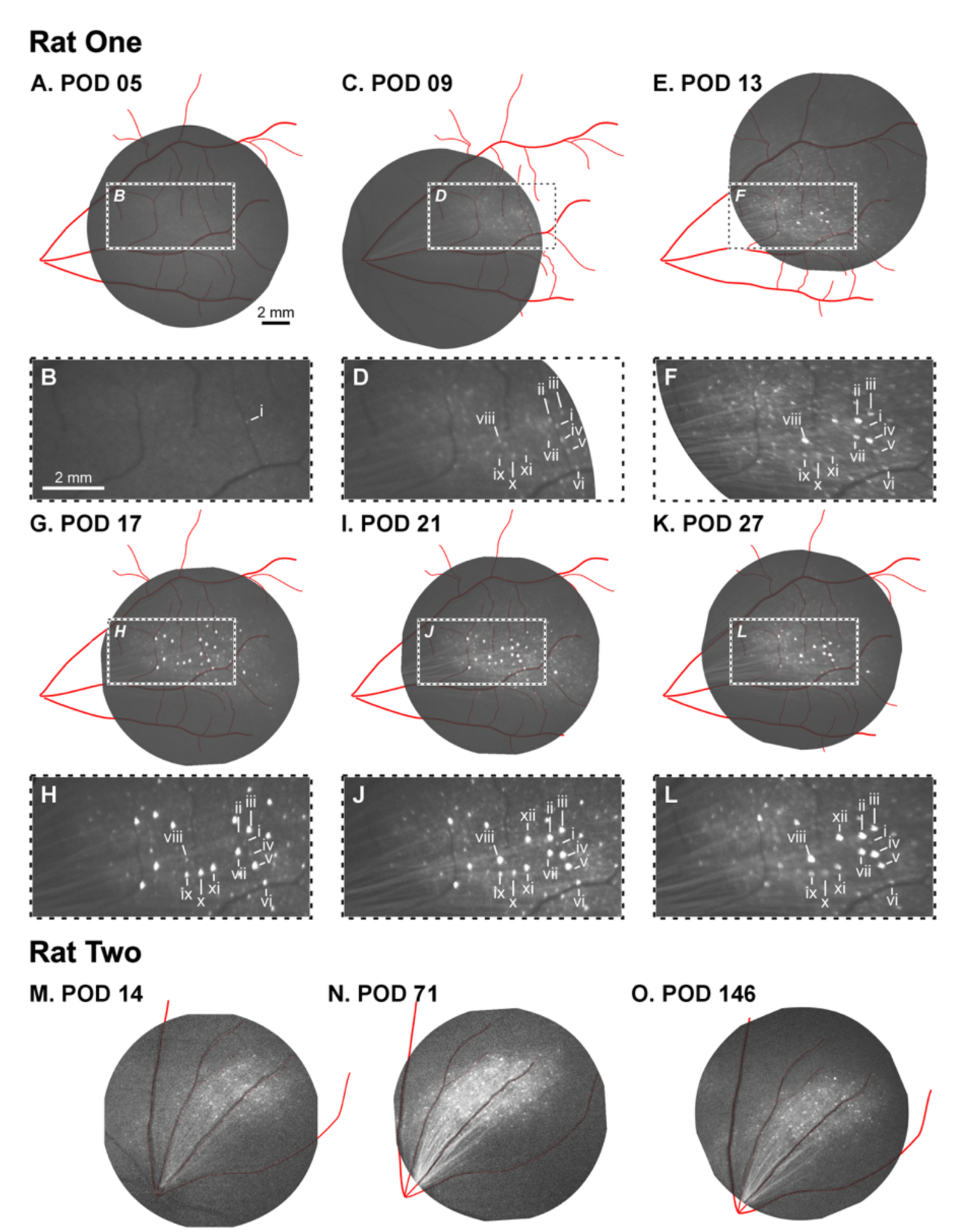
Identification of transduced RGCs using *in vivo* fluorescence fundoscopy. Fluorescence fundus images are shown from two example rats. The schematic for rat one illustrates the left eye at post-operative days (POD) (**A**) 5, (**C**) 9, (**E**) 13, (**G**) 17, (**I**) 21, and (**K**) 27. Red lines depict the intraocular vasculature used to align fundus images across days. Insets (**B, D, F, H, J,** and **L**) show magnified views of regions with prominent labeling, as indicated by the dashed white rectangles (**A, C, E, G, I,** and **K,** respectively). Examples of transduced RGCs are indicated with Roman numerals (**i-xii**), several of which are present across multiple imaging timepoints. The schematic for Rat Two (**M-O**) illustrates an example of expression longevity starting from POD 14 through POD 146.

We next determined the extent to which fluorescence fundus images accurately revealed the location and density of labeling in the retina. To achieve this, we compared fundus images to post-mortem confocal microscopy images of the same retinas after histological amplification of the mCitrine signal (**Fig. 3**). Although dendritic processes could not be identified from fundus images, the locations of fluorescently labeled somas were strongly correlated between fundus and confocal images (**Fig. 3E-H**). This indicates that fluorescence fundoscopy is effective for identifying the location and density of transduced cell bodies within the retina *in vivo (Nanjappa et al., 2022; Zheng et al., 2024)*.

**Figure 3.**
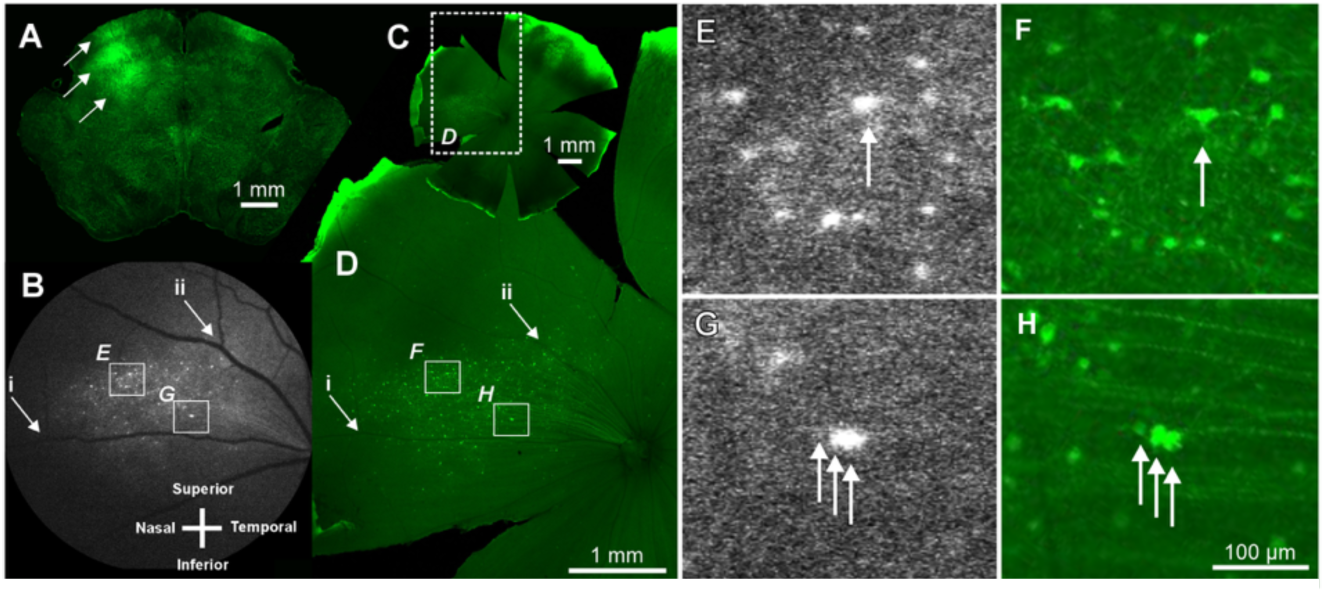
Comparison of fluorescence fundoscopy and immunofluorescence confocal imaging. (**A**) Injection of AAV2retro-CAG-ReaChR-mCitrine into the left SC resulted in labeling across tectal layers, as indicated by the arrows. (**B**) This injection resulted in fluorescence fundoscopic detection of mCitrine-positive RGC labeling within the nasal quadrant of the right eye. (**C**) A low-magnification confocal photomicrograph of the eyecup following anti-GFP immunofluorescent amplification. The dashed box in C delineates the region shown in D. (**D**) A confocal photomicrograph shown at the estimated scale as the fluorescence fundus image in B. Alignment of these images was achieved using the major blood vessels. Roman numerals **i** and **ii** identify examples of vessel bifurcation used to match the images from fluorescence fundoscopy with those from confocal imaging. (**E-H**) Examples of retrogradely transduced RGCs are depicted (arrows) for aligned fluorescence fundoscopy images (**E, G**) and confocal images (**F, H**). Boxes in B reflect regions illustrated in E and G, and boxes in D reflect regions illustrated in F and H.

### Using MEAs to measure visual responses from transduced cells

We next sought to measure the visual responses of SC-projecting RGCs using a large-scale MEA. Segments of the retina with labeled RGCs were placed on the MEA (**Fig. 4A, B**; see Methods). These segments were identified by nearby vasculature, which was clearly visible in the fundus images (**Fig. 4A**) and in the isolated retina viewed under infrared (IR) illumination with the use of IR converters on a stereomicroscope (not shown). Following placement of the retina onto the MEA, visual stimuli were presented to measure RGC receptive fields (RFs) and other response properties (see Methods).

**Figure 4.**
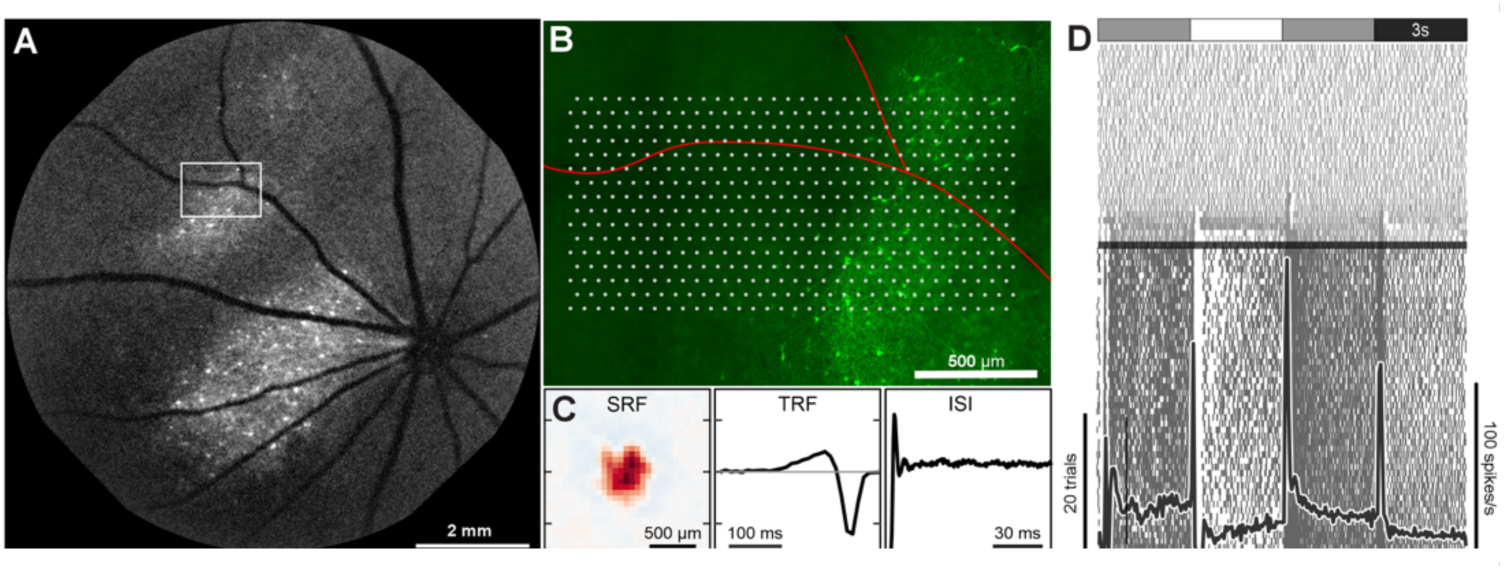
Measuring visual responses of transduced RGCs using the MEA. (**A**) Example case of fluorescence fundoscopy illustrating retinotectal RGC labeling and the nearby vasculature. The white box illustrates the approximate location of the retinal piece that was placed onto the MEA. (**B**) Fluorescence photomicrograph of the retinal piece with retrogradely labeled retinotectal RGCs (green cells), electrodes (white dots) of the MEA, and blood vessels (red lines). (**C**) Spatial receptive field (SRF), temporal receptive field (TRF), and inter-spike interval distribution (ISI) of an example RGC over the MEA. (**D**) Raster in response to full-field light steps switching from gray-to-white-to-gray-to-black every 3 s (top). Drug wash-on is indicated by the black horizontal bar. Prior to wash-on (lower portion of raster), the cell robustly responded to light steps: superimposed black trace is the peri-stimulus time histogram of the spike activity prior to drug wash-on. Shortly after the drug wash-on (to block photoreceptor signals), the spiking becomes uncorrelated with the stimulus (upper portion of raster).

Following previous work, we used checkerboard noise, drifting gratings, and full-field light steps to distinguish RGC types (Ravi et al., 2018). Checkerboard noise was used to estimate the spatial and temporal components of RGC RFs by computing the spike-triggered average (STA). The inter-spike interval distribution (ISI) measured during the checkerboard noise was also used for functionally distinguishing the RGCs (**Fig. 4C**). Drifting gratings allowed identification of direction-selective and orientation-selective RGCs (Yao et al., 2018). Finally, full-field light steps that transitioned from gray-to-white-to-gray-to-black every 3s (**Fig. 4D**) were used for distinguishing ON, OFF, and ON-OFF RGCs. Full-field light steps were also used for measuring the effects of drugs to block photoreceptor-mediated light responses (**Fig. 4D**; see Methods). Following the administration of the drugs, RGC spiking activity exhibited no modulation by the light steps (raster rows in the top half of **Fig. 4D**), indicating that the photoreceptor-mediated responses were blocked. This allowed isolating ReaChR-mediated responses in RGCs labeled by the AAV2retro injections into the SC.

### Identifying ReaChR-expressing RGCs on the MEA

To match the visual responses of RGCs recorded on the MEA to ReaChR-expressing cells (**Fig. 4B**), we used an approach called phototagging. Following the pharmacological block of photoreceptor-mediated responses (**Fig. 4D**), bright full-field flashes from a 565 nm LED were presented to drive ReaChR responses (**Fig. 5A**). These flashes produced a short latency increase in the spiking of some RGCs over the MEA with longer flashes driving more spikes (e.g., **Fig. 5A**; left column). In contrast, most RGCs showed no significant changes in spike rate in response to the flashes (e.g., **Fig. 5A**; right column). To distinguish these two populations of RGCs, we developed an automated classifier (see Methods). Briefly, the classifier was based on distinguishing cells with a flat versus increasing intensity-response relationship to the LED flashes (**Supp.** Fig. 1). RGCs lacking ReaChR expression exhibited a Gaussian distribution of intensity-response profiles near zero (flat), while RGCs with ReaChR expression were outliers from this distribution (**Fig. 5B**; top). As a control, when this analysis was applied to retinas from rats that did not receive SC injections (and had no ReaChR expression), the false positive rate of RGCs exceeding the threshold of four standard deviations was zero (**Fig. 5B**; bottom).

**Figure 5.**
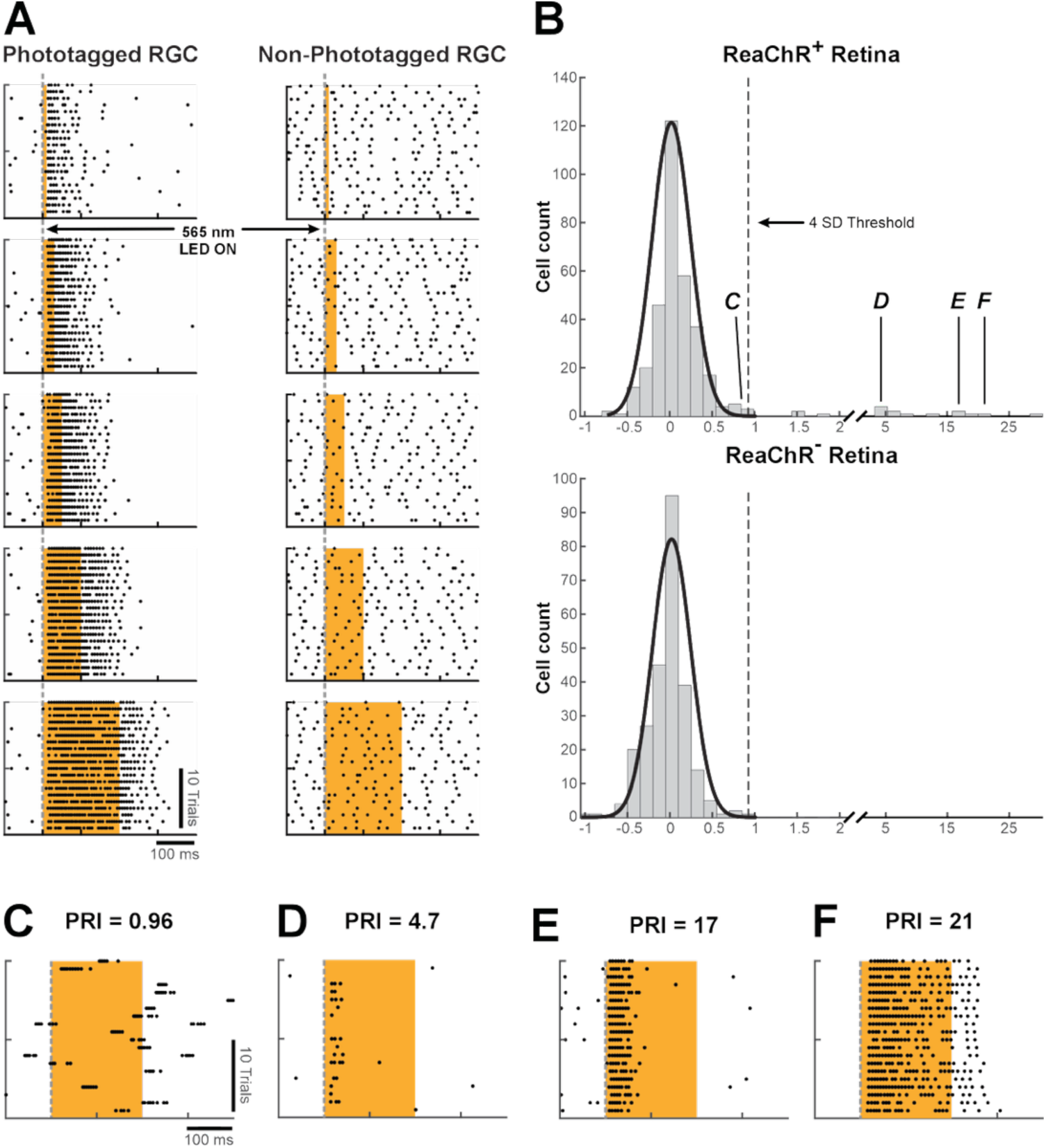
Identification of phototagged ReaChR-expressing cells. (**A**) Spike rasters to the presentation of 565 nm LED flashes (orange bars) for two example RGCs from the same experiment (left column: phototagged or ReaChR-positive; right column: non-phototagged or ReaChR-negative). Each row depicts a longer duration flash (from top to bottom): 10 ms (generated 1.97^13^ photons/mm^2^); 30 ms (5.90^13^ photons/mm^2^); 50 ms (9.83^13^ photons/mm^2^); 100 ms (1.97^14^ photons/mm^2^); 200 ms (3.93^14^ photons/mm^2^). (**B**) Distributions of Phototagged Response Index (PRI) for a retina with cells containing ReaChR (top) and a control retina with cells lacking ReaChR expression (bottom). The two example retinas were presented with the same flash strengths and durations. Labels ***C-F*** indicate example cells shown below. (**C**) Spike raster for a cell immediately below the PRI threshold of 4 SD, indicating that the cell was not photosensitive to 565 nm LED flashes (3.93^14^ photons/mm^2^). (**D-F**) Spike rasters for weakly (**D**), moderately (**E**), and strongly (**F**) ReaChR-activated RGCs.

Importantly, the visual stimuli used to measure photoreceptor-mediated responses were presented at much dimmer intensities than the LED flashes needed to drive ReaChR responses (∼10,000 photoisomerizations per rod per second for photoreceptor-mediated responses compared with >25,000,000 photoisomerizations per rod per second for ReaChR responses). Thus, photoreceptor-mediated responses were not contaminated by ReaChR stimulation, and ReaChR-mediated responses were not contaminated by photoreceptor-mediated responses because of the pharmacological block.

Once the population of ReaChR-expressing RGCs was identified, their photoreceptor-mediated visual responses were analyzed (**Fig. 6**). In an example experiment, ReaChR-expressing RGCs were distributed along the lower portion of the MEA (**Fig. 6A**; left). These RGCs displayed a range of response properties, including diverse temporal RFs (**Fig. 6A**, center-left), ISI distributions (**Fig. 6A**, center-right), and contrast response functions (CRFs; **Fig. 6A**; right). RGCs were tracked across stimulus and drug conditions by their electrical images (EIs; **Fig. 6B-E**; top row). EIs are the spike-triggered average electrical activity over the MEA and serve as a “footprint” for tracking individual cells across conditions (Field et al., 2009; Yao et al., 2018). This allowed matching the ReaChR-expressing, fluorescently labeled RGCs to their visual response properties.

**Figure 6.**
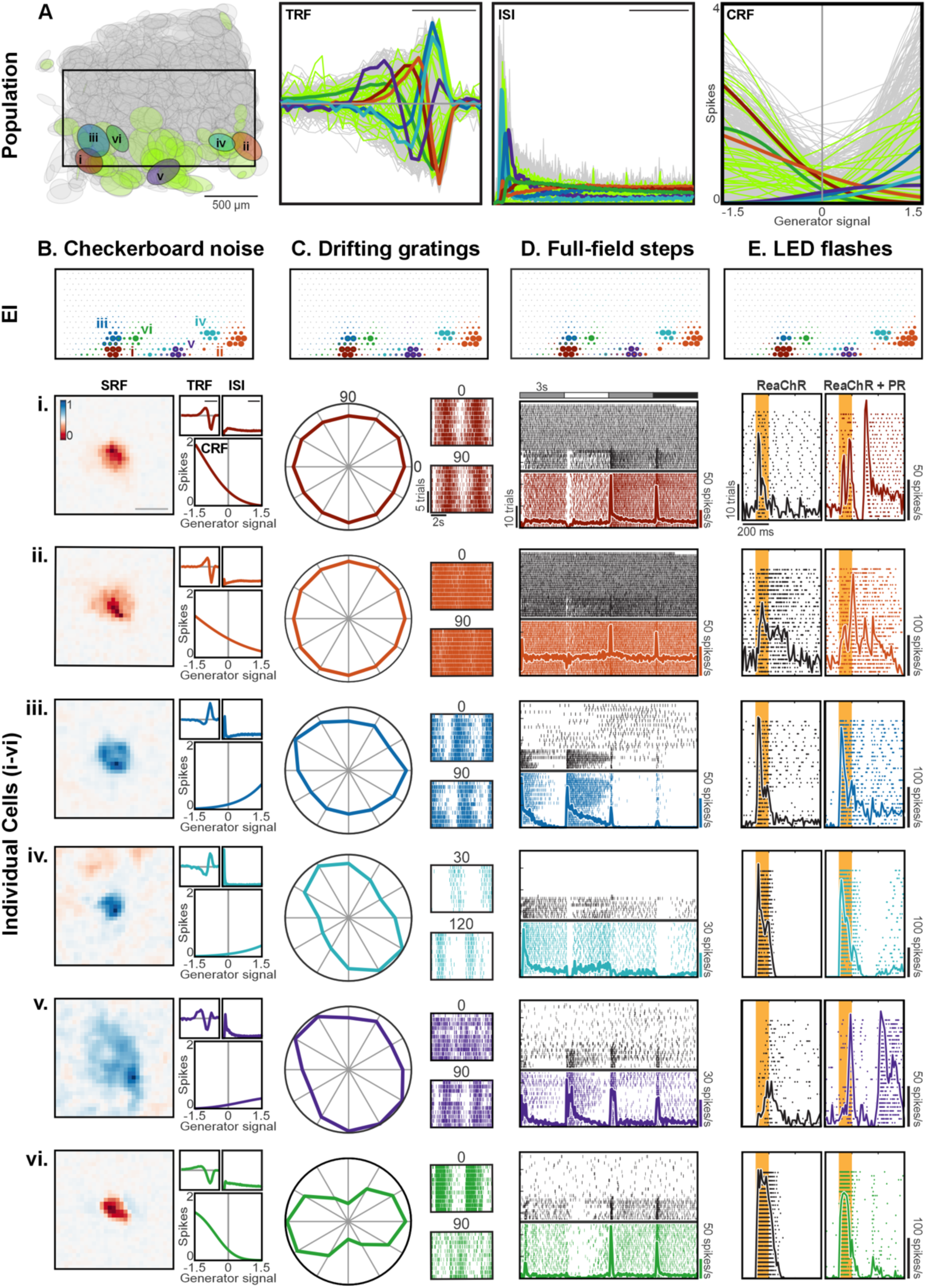
Diverse visual response properties of phototagged RGCs. (**A**) *Left*: Spatial receptive field (SRF) fits for all RGCs (N=1175) in one experiment. Each ellipse is the 1-SD contour of a 2-dimensional Gaussian fit. ReaChR-negative RGCs are gray, ReaChR-positive RGCs are green, and other colored ellipses highlight example ReaChR-positive RGCs shown in panels B-E. *Left-middle:* Temporal receptive fields (TRFs) shown for the same cells in A; scale bar is 100 ms. *Right-middle:* Inter-spike interval (ISI) distributions shown for the same cells in A; scale bar is 30 ms. *Right*: Contrast response functions (CRFs) shown for the same cells in A. **(B-E)** Visual response properties and electrical images (EIs) of six example ReaChR-positive RGCs are shown across four stimulus conditions. EIs for all six example cells are shown together for each stimulus condition (top). (**B**) SRF (scale bar = 500 µm), TRF (scale bar = 100 ms), ISI (scale bar = 30 ms), and CRF were calculated from the checkerboard noise stimulus. (**C**) Responses to drifting gratings presented in 12 directions are shown in polar plots (left) and spike rasters for two indicated directions (right). Each polar plot is normalized to the maximum spike count. (**D**) Spike rasters (ticks) and peri-stimulus time histograms (bold colored lines) are shown for full-field light steps, during which the stimulus repeatedly transitioned from gray-to-white-to-gray-to-black every 3 s (top). Photoreceptor-mediated responses were measured first (colored portion of raster), then drug cocktail was applied (black horizontal line) to block synaptic transmission between photoreceptors and bipolar cells (see Methods). (**E**) Left column shows ReaChR responses to a 100 ms 565 nm LED flash (orange bar) during pharmacological block of photoreceptor-mediated responses. Right column shows ReaChR and photoreceptor-driven (PR) responses obtained after drug wash out. Flash strength was ∼10^14^ photons/mm^2^.

For six example ReaChR-expressing RGCs, responses to checkerboard noise, drifting gratings, full-field light steps, and the 565 nm LED flashes are shown (**Fig. 6B-E**). Example cells *i-iv* exhibited approximately Gaussian spatial RFs (**Fig. 6B**; left). However, these cells differed in their spatial RF sizes, response polarities, temporal integrations, and contrast response functions (**Fig. 6B**; right). In general, these cells were neither direction-selective nor orientation-selective, though cell *iv* exhibited weak orientation selectivity (**Fig. 6C**; Orientation Selectivity Index (OSI) = 0.29; see Methods). Cell *v* exhibited a diffuse, non-Gaussian spatial RF and clear ON-OFF responses to the full-field light steps. Cell *vi* exhibited clear orientation selectivity (**Fig. 6C**; OSI = 0.54) that was produced by a spatial RF consisting of an OFF center and two flanking ON sub-regions reminiscent of an orientation-selective simple cell in primary visual cortex (Hubel and Wiesel, 1962). We also observed ReaChR-expressing RGCs that were direction-selective in many (but not all) of these experiments (**Supp.** Fig. 2). All six example RGCs exhibited short-latency ReaChR-driven responses to the bright LED flashes (**Fig. 6E**; left) and longer latency photoreceptor-driven responses upon washing out the pharmacological blockade of photoreceptor output (**Fig. 6E**; right).

### Matching visual responses to morphology

The goal of fluorescently labeling RGCs and identifying their visual responses via phototagging was to match cell morphology to function. To achieve this, the retina was imaged with an epifluorescent microscope following the assessment of photoreceptor- and ReaChR-mediated visual responses (**Fig. 6**) and prior to removing it from the MEA (**Fig. 7A, B**). This allowed visualization of fluorescently labeled RGCs and their relative positions on the MEA. Subsequently, the retina was drop-fixed in 4% PFA, and the viral-mediated fluorescent signal was amplified by immunofluorescence (see Methods). Additionally, choline acetyltransferase (ChAT) antibody was used to landmark the inner plexiform layer (IPL)(Manookin et al., 2008). Confocal images of the piece of retina were then acquired (**Fig. 7C**), providing higher resolution images for distinguishing RGC morphologies. Based on anatomical landmarks, including retinal vasculature and the locations of fluorescent RGCs, we approximated the MEA borders on the confocal images (**Fig. 7C**, solid black rectangle) from the live images (**Fig. 7A**).

**Figure 7:**
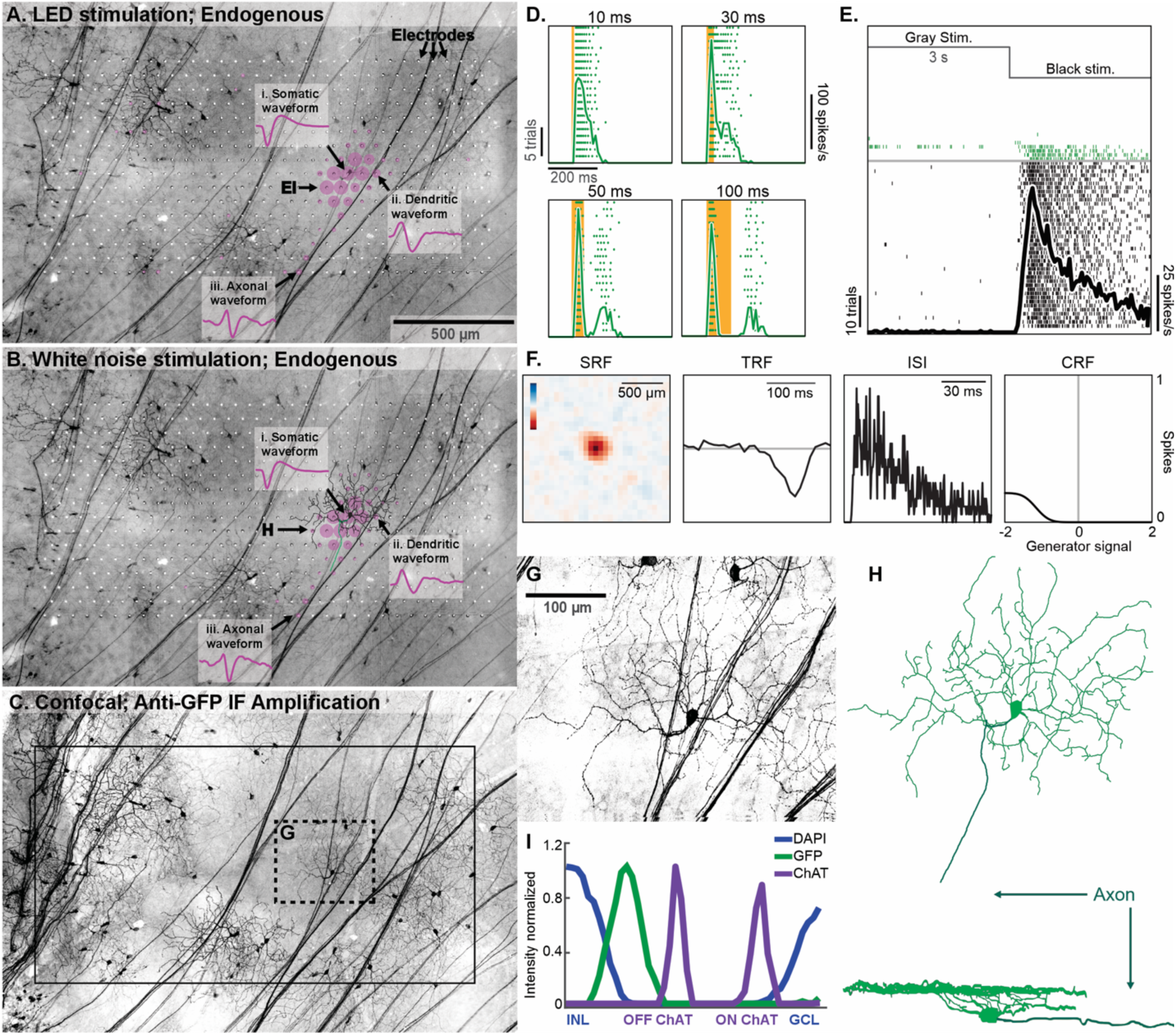
Linking physiology with morphology. All plots depict data from one example RGC. **(A)** Photomicrograph of the *ex vivo* retina on the array with the EI (pink) of one RGC superimposed. EI was computed from ReaChR responses driven by the 565 nm LED with drugs to block the photoreceptor-mediated response. Spike waveforms from 3 electrodes are shown: one near the soma of the RGC, one near the distal dendrites, and one along the axon. Waveform shapes and polarities are consistent with expectations for somatic, dendritic, and axonal signals. **(B)** EI obtained during checkerboard noise stimulation, depicted for the same example RGC and photomicrograph as shown in A. EI shape and waveforms match those during LED stimulation in A. A post hoc reconstruction from confocal images of the example RGC (also illustrated in H) is overlaid on the EI and demonstrates remarkable congruence between the EI footprint and the cell’s morphological features. **(C)** A 4x confocal photomicrograph of the example retina following immunofluorescent (IF) amplification. The solid black box indicates the array borders, and the dashed box indicates the region imaged at higher magnification in G. **(D)** Spiking activity evoked by 10, 30, 50, and 100 ms 565 nm LED flashes following drug wash-on, indicating that the RGC highlighted in A and B is phototagged and thus projects to the SC. **(E)** Before the drug wash-on (gray horizontal line), the example RGC strongly responded to a light step from gray to black (spikes indicated by black ticks). The subsequent rapid loss of sensitivity following drug application confirms that the optogenetic responses in D are not mediated via photoreceptor activation. **(F)** Shown are the spatial receptive field (SRF), temporal receptive field (TRF), inter-spike interval (ISI), and contrast response function (CRF) derived from checkerboard noise stimuli. **(G)** 60x confocal photomicrograph of the example RGC was reconstructed from a z-stack (70 planes). **(H)** Morphological reconstruction of the example cell was derived from high-magnification confocal images in G, shown from a top-down view (top) and a side view (bottom), with the axon shown in dark green. **(I)** Analysis of the mean fluorescence intensity across the z-stack indicates that the cell’s dendritic processes stratify within the OFF sub-lamina of the IPL.

ReaChR-positive RGCs were then identified after spike sorting (**Fig. 7D**). The EIs of these ReaChR-positive cells were superimposed over the images of the fluorescent RGCs on the MEA (**Fig. 7A**) and mapped to other stimulus conditions, such as checkerboard noise (**Fig. 7B**). This allowed matching the visual responses from individual RGCs across different stimuli and to the receptive field properties estimated from the STA (**Fig. 7E-F**). Then, the confocal images of ReaChR-positive RGCs (**Fig. 7C, G**) were analyzed to determine the detailed dendritic morphologies (**Fig. 7H**) and stratification depths in the inner plexiform layer (**Fig. 7I**). Together, these steps allowed matching the receptive fields and other visual response properties of labeled RGCs to their morphologies.

Additional confirmation of matches between morphology and physiology was obtained by comparing the EI spike waveforms on individual electrodes and their proximity to different neuronal compartments. From an extracellular electrode near the soma, a spike produces a downward followed by an upward voltage deflection (Litke et al., 2004; Petrusca et al., 2007) (**Fig. 7Ai**). Electrodes near dendrites exhibit a sign-inverted signal relative to the soma, and this was observed on electrodes near dendrites (**Fig. 7Aii**). Finally, traveling axonal spike waveforms are triphasic, and this waveform shape was observed on electrodes next to the axon (**Fig. 7Aiii**). These waveforms were consistent across EIs from ReaChR- and photoreceptor-driven spikes (**Fig. 7A-B i-iii**), indicating that the EI did not depend strongly on whether spikes were driven by ReaChR or synaptic input. Collectively, these observations allowed us to reliably match the physiology and morphology of the labeled RGCs.

Finally, two factors yielded more confidence in this match between visual responses and morphology. First, the STA of the example RGC exhibited an OFF-center spatial RF and a relatively sustained response from the primarily monophasic temporal RF (**Fig. 7F**). The morphology of the matched RGC was consistent with these visual responses, given prior knowledge of retinal anatomy: the dendrites stratified in the outer portion of the IPL (**Fig. 7I**), outer to the ‘OFF’ ChAT band, where OFF-sustained RGCs send their dendrites (Baden et al., 2013; Borghuis et al., 2013; Bae et al., 2018). Second, the spatial extent of the dendritic field Ωwas also well-matched to the size of the spatial RF. The dendritic field area was quantified from a convex hull (**Fig. 7H**; A = 9.2 × 10⁴ µm²). The spatial RF area was estimated from a two-dimensional difference-of-Gaussians fit to the STA (**Supp.** Fig. 3; A₁ = 1.77 × 10⁴ µm²; A₂ = 7.06 × 10⁴ µm²). Prior work indicates that the dendritic field spans approximately a two-standard-deviation contour of the RF center (Baden et al., 2016). The radius of the dendritic field area for the example cell was quite close to that of the radius of the Gaussian fit at two standard deviations (160 µm vs. 150 µm, respectively). These comparisons, as well as comparisons to the visual responses of other nearby RGCs with similar Eis, indicated a reliable matching procedure (**Supp.** Fig. 4).

Based on somatic size, dendritic field area, and branching complexity, the example RGC closely resembles morphological cell group C, as described previously in the rat retina (Sun et al., 2002). However, current catalogues of rat RGC types are likely incomplete, given that they consist of ∼20 RGC types, while mouse catalogues contain ∼45 RGC types. Thus, we also compared the features of this example RGC to published mouse catalogues. The sustained OFF response along with the soma size and dendritic field morphology was most consistent with the 2aw type^1^ (Bae et al., 2018; Goetz et al., 2022), though the 1no and 3o types are also possibilities: it remains unclear how to precisely match RGC types between mouse and rat.

### Using RGC mosaics to identify retinotectal RGC types

In some experiments, the distribution of transduced RGCs was dense, which made it challenging to trace individual neurons (**Fig. 7A** compared with **Fig. 8A**). In these cases, we were frequently able to use the organization of RGCs into mosaics that tile the retina to identify irreducible types that project to the SC. For example, we functionally classified RGCs by their visual responses (e.g., RF properties, CRFs, and ISI distributions) using methods described previously (**Fig. 8**)(Ravi et al., 2018). Mosaics of RFs would emerge from this classification, which were apparent when comparing the spatial arrangement of all RFs (**Fig. 8B**) to just those RGCs that exhibited stereotyped response properties (**Fig. 8C**). In this example, the OFF brisk-transient RGC type in the rat retina is shown (Crook et al., 2008; Krieger et al., 2017; Ravi et al., 2018). Most of these RGCs were ReaChR-positive (23 of 39; **Fig. 8C**, green vs. gray ellipses for ReaChR-positive and negative RGCs, respectively) and nearly all were positive on the side of the MEA that had labeled cells. Thus, in addition to matching individual cells, the projection targeting with phototagging approach enables entire populations of irreducible cell types to be identified as projecting to the SC in densely labeled areas of the retina.

**Figure 8:**
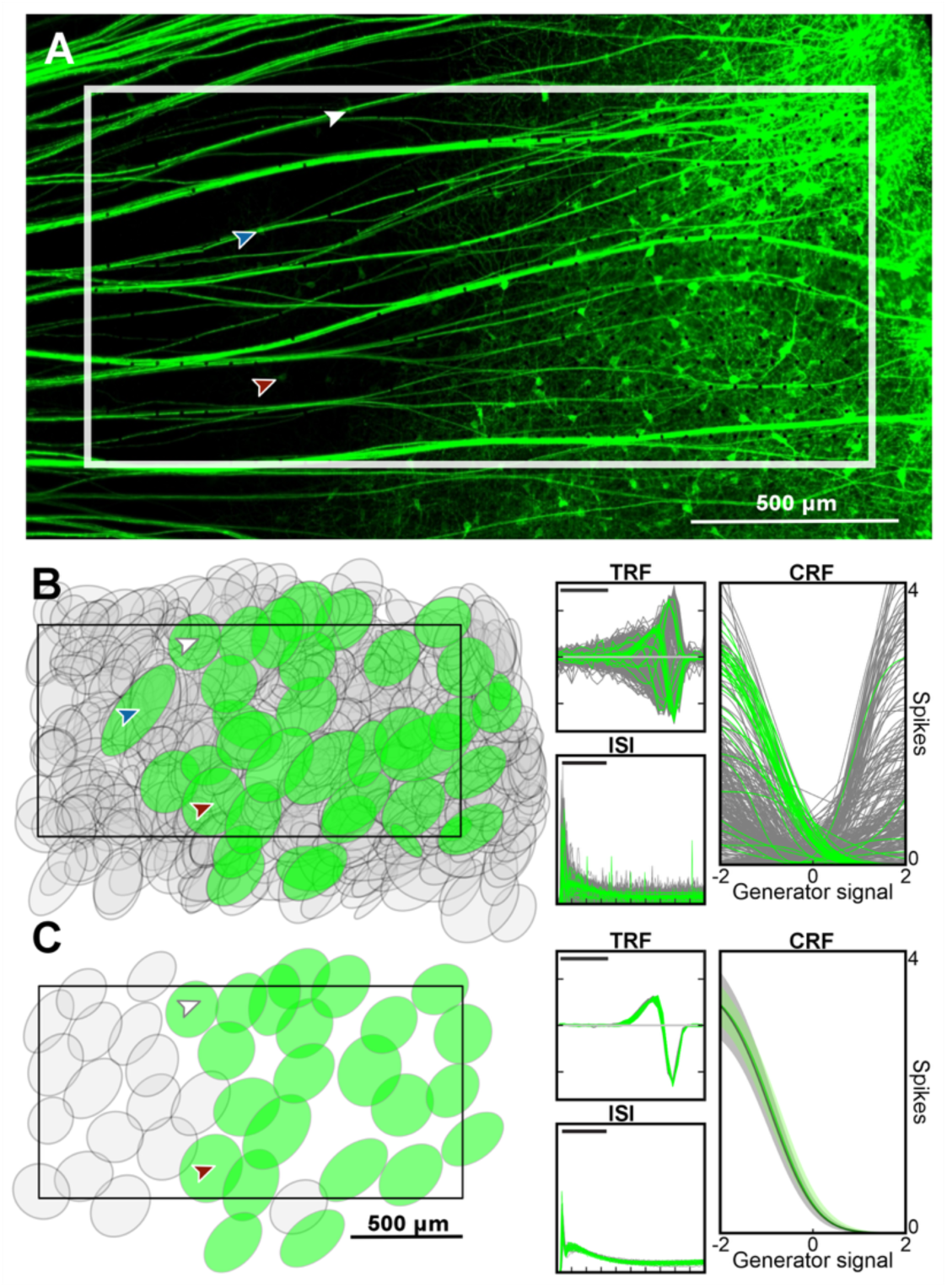
Matching ReaChR-positive RGCs to functional types using mosaics. **(A)** Photomicrograph of the *ex vivo* retina on the MEA (white rectangle). **(B)** Left plot shows spatial receptive field fits for all identified RGCs (n = 546) over the MEA in A. Each ellipse is the 1-SD contour of a 2-dimensional Gaussian fit. Gray ellipses are ReaChR-negative RGCs (n = 508), green ellipses are ReaChR-positive RGCs (n = 38). The rectangle shows the outline of MEA. Arrowheads match RGCs in B to faintly labeled RGCs in A. Right plots show temporal receptive fields (TRFs) (scale bar = 100 ms), inter-spike interval distributions (ISI) (scale bar = 30 ms), and contrast response functions (CRF) derived from the checkerboard noise stimulus. **(C)** Same as B, but for the OFF brisk-transient RGCs (n = 40). The average CRF is shown (phototagged in green and non-phototagged in black) across the identified OFF brisk transient RGCs, with the shaded area indicating the 1-SD boundary.

## DISCUSSION

This study refined an approach, projection targeting with phototagging, and demonstrated that it is useful for interrogating and matching the structures, visual responses, and central projections of RGC types. By delivering light-sensitive opsins to RGCs via retrograde virus injections in rat SC and pairing this with large-scale MEA recordings, we showed it is possible to phototag individual RGCs according to their projection target, characterize their light-evoked response properties, reconstruct their morphologies, and identify their axonal projections—all without genetically modified animals. Projection targeting with phototagging is informative, high-throughput, and scalable, and, due to its reliance on viral vectors rather than transgenic lines, it offers a versatile platform for mapping the early visual pathway across diverse mammalian species.

### Comparisons to prior approaches

#### Conventional methods for morphological identification

Early morphological studies of RGCs relied on injections of non-viral retrograde tracers into retinorecipient structures or the use of direct, intracellular fills (see references for examples in primate (Dacey et al., 2003), cat (Wassle and Illing, 1980), rabbit (Vaney et al., 1981), squirrel (Rivera and Lugo, 1998), and rat (Linden and Perry, 1983)). However, these methods have significant limitations. Conventional tracers like Wheat Germ Agglutinin or Cholera toxin subunit B rarely provide complete intracellular fills, making it challenging to reconstruct detailed RGC morphology. Furthermore, these tracers suffer from fibers-of-passage uptake at the injection site. Conventional intracellular fills, while providing detailed morphological insights, are labor intensive and can be biased toward numerically dominant cell types. Combining retrograde tracer with intracellular injections can enhance morphological detail, but such methods remain time-consuming and technically demanding.

In comparison, projection targeting with phototagging relies on injecting AAV2retro into retinorecipient structures. The virus is taken up at the RGC axon terminals in the SC and retrogradely transported to the nucleus where virally conferred fluorescent proteins are expressed. We found that this expression resulted in a putatively complete intracellular fill of transduced RGCs, revealing remarkable morphological details (**Supp.** Fig. 5). Compared with conventional tracers, which often lead to nonspecific fibers-of-passage uptake, AAV vectors selectively transduce neurons at their terminals ensuring more precise and targeted labeling (Chamberlin et al., 1998).

#### Previous approaches for matching RGC morphology and axonal projections to visual responses

Several approaches have utilized genetically modified mouse lines to track the projection patterns of RGCs and match visual responses to cellular morphology (Huberman et al., 2008; Yonehara et al., 2008; Huberman et al., 2009; Kim et al., 2010). These approaches have proven powerful for understanding the mouse visual system but have raised questions about the extent to which the mouse is similar or different from other mammals (particularly primates) in terms of RGC diversity and projection patterns. Answering these questions will likely require the development and deployment of approaches that do not rely on genetically modified animals.

In comparison to projection targeting with phototagging, perhaps the most similar previous approach involves rhodamine dextran injections into retinorecipient areas followed by single unit recordings from filled cells (Dacey et al., 2003). That approach allows individual RGCs to be targeted for sharp electrode or patch-clamp experiments while also providing complete fills of dendritic morphologies following exposure to bright light. However, the throughput of the electrophysiological measurements and label uptake is much lower than the approach described here. Another method involves using transsynaptic rabies-based tools to deliver exogenous genes for labeling and/or manipulating neurons. This approach has been used in mouse to generate catalogs of RGCs with matched morphologies and visual responses, but it has not been combined with large-scale MEA experiments (Reinhard et al., 2019).

To match visual responses to projections, antidromic electrical stimulation has also been employed, but this approach does not reveal the morphology of the RGCs (Schiller and Malpeli, 1977). Another approach has been to unselectively patch-clamp individual RGCs, measure visual responses and/or synaptic inputs, and then dye fill cells for morphological reconstructions. However, this approach is low-throughput and does not, in general, provide information about the locations to which RGCs project. Further, this approach does not typically provide clear classification schemes because it does not reveal the presence (or absence) of mosaics, a hallmark of RGC types.

An alternative to central injections into retino-recipient areas with retrograde viral labeling is to inject viruses intravitreally. This approach has a few issues. First, the inner-limiting membrane can limit viral access to the retina, preventing RGC labeling (Godat et al., 2022). Second, this approach does not readily indicate which RGCs project to different brain areas. Third, as noted below, it can result in pronounced immune responses (Godat et al., 2022).

High-throughput matching of cellular morphology with physiological responses is a general challenge in neuroscience. In the retina, anatomical features reduce some barriers to achieving this goal. In particular, the planar arrangement of RGCs facilitates the use of planar electrode arrays. This planar arrangement also allows EIs to be calculated that reflect the somatic location, dendritic field position and axonal trajectory toward the optic nerve head (Litke et al., 2004; Petrusca et al., 2007; Li et al., 2015). These features of the EIs have proven useful for tracking RGCs across stimulus and drug conditions in previous studies (Field et al., 2009; Yao et al., 2018). Here we use the EI to reliably match ReaChR-mediated to photoreceptor-mediated visual responses of virally labeled RGCs, thus linking the projections, physiologies, and morphologies of individual cells. However, using the EI and mCitrine expression alone, without phototagging (ReaChR expression), it would have proven challenging in some cases to reliably match the visual responses to the morphologies of cells. Furthermore, this matching primarily works in regions where the labeling is relatively sparse: it becomes more challenging to trace individual cell morphologies when the labeling is dense. To overcome this challenge, we are investigating techniques such as expansion microscopy and machine learning algorithms to improve neurite tracing and cell identification in densely labeled areas of the retina.

### Technical considerations

There are several technical issues to consider when applying the virally mediated retrograde phototagging approach we have described.

#### Viral capsid tropism, promoter interactions, and expression

The choice of viral capsid and promoter is important. We used AAV2retro due to its high propensity for efficient retrograde transport and transduction efficiency to projection neurons (Tervo et al., 2016). However, interactions between the viral capsid and the promoter used to drive transgene expression may influence transduction outcomes (Bohlen et al., 2020). The extent to which capsid-promoter interactions may bias the labeling of specific RGC types remains an open question. In the current study, we used the CAG promoter, which targets neuronal and non-neuronal cells. Combining traditional dye injections with viral tracers may be useful for better understanding these biases. In addition, future work should examine the efficacy of other promoters in combination with AAV2retro for labeling RGCs. Comparisons should be made between the number and diversity of RGCs labeled when different promoters are used.

#### Immune responses

Immune responses to viral vectors are another critical consideration. In the current study, we did not observe an obvious immune response following injections into the SC. This suggests that central administration may be an immunologically safer method for delivering transgenes to RGCs compared to intravitreal injections, which can provoke stronger immune reactions in the eye (Godat et al., 2022).

#### Variability in viral uptake and expression patterns

In this study, the greatest variability in uptake and expression was primarily attributed to injection placement. Injections concentrated in the deep and intermediate SC resulted in a focused region of retrogradely labeled RGCs, while more superficial SC injections resulted in comparatively diffuse labeling of RGCs (**Fig. 1H**). We hypothesize that this may result from how the virus diffuses, with more diffusion in superficial layers than in deeper layers. Lateral SC injections resulted in superior RGC labeling (**Fig. 1I**), more medially placed injections resulted in more inferior retinal labeling (**Fig. 1I**), rostral SC injections produced temporal RGC labeling (**Fig. 1J, K**), and caudal SC injections resulted in nasal RGC labeling (**Fig. 1J, K**). This systematic topography opens the potential to study whether there are functional or morphological differences in the RGCs that project to different regions of SC.

### Future directions

Recent advances in the understanding of the mammalian retina have highlighted the diversity of RGC types. The mouse retina has ∼45 RGC types (Martersteck et al., 2017), while primates have at least ∼20 types (Dacey et al., 2003; Yamada et al., 2005; Peng et al., 2019; Kling et al., 2024). These RGC types project to diverse brain targets, possibly as many as ∼50 areas (Martersteck et al., 2017). However, much of this knowledge is based on findings from the mouse with comparatively less known about RGC morphology, physiology, and projections in primate, cat, rabbit and other species. To build a more complete understanding of how RGCs shape visual processing and to provide controlled, replicable comparisons of retinal function across species, it would help to determine the morphological and functional diversity of RGC types and their projection patterns using methods that remain as constant as possible across studies. The approach we describe here, projection targeting with phototagging, could serve this effort. Furthermore, this approach would allow for cells to be sorted by fluorescence, permitting a comparison of RGC transcriptomics with projection patterns. This knowledge would provide greater insight into the relationship between gene expression profiles and the structure-function of evolutionarily conserved (or divergent) visual pathways(Baden et al., 2020; Hahn et al., 2023). This scope and scale are necessary for understanding how evolution has modified the mammalian visual system to meet the challenges of different ecological niches.

## SUPPLEMENTAL FIGURES

**Supplemental Figure 1.**
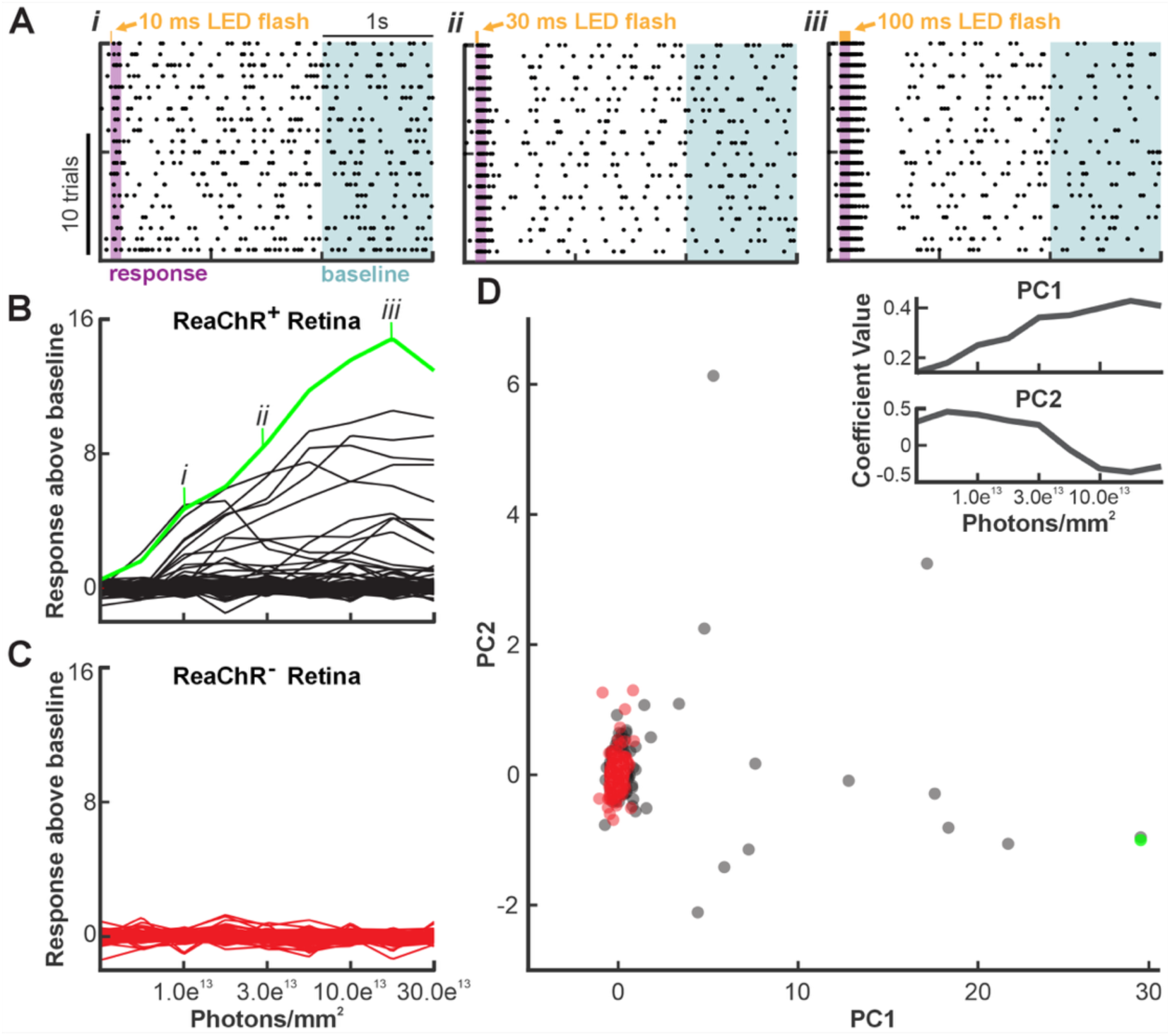
Identification of ReaChR-positive RGCs. (**A**) Spike rasters to three stimulus conditions that increasingly drive the ReaChR response in an example RGC. Flash duration (orange bars at tops of panels) increases from left to right: *i* = 10 ms (1.97^13^ photons/mm^2^), *ii* = 30 ms (5.90^13^ photons/mm^2^), and *iii* = 100 ms flashes (1.97^14^ photons/mm^2^). Flashes were delivered with a 565 nm LED while applying a drug cocktail to block the photoreceptor-mediated response (see Methods). At each flash strength, the ReaChR response (e.g., response above baseline in B and C) is calculated as the average spike rate during the 100 ms period following the flash onset (purple bars) minus the average baseline spike rate during the 2-3 s period following each flash (blue bars). Averages were calculated over trials. (**B**) Responses as a function of flash strength (total number of photons per flash) for all RGCs in a ReaChR-positive retina. The example phototagged RGC from A is shown in green. (**C**) Responses as a function of flash strength in a ReaChR-negative retina. (**D**) Weights associated with the first and second principal components (PC) derived from principal component analysis applied to the response vs. flash strength functions shown in B (gray) and C (red); example cell from A shown in green. Insets show the 1^st^ and 2^nd^ PCs. PC1 and PC2 captured 92.2% and 3.7% of the variance, respectively. PC1 captured most of the variance in the data and provided an effective axis for distinguishing ReaChR-positive cells (the outliers) from ReaChR-negative cells (clusters). The weights along PC1 form the Phototagged Response Index (PRI) in Figure 5.

**Supplemental Figure 2.**
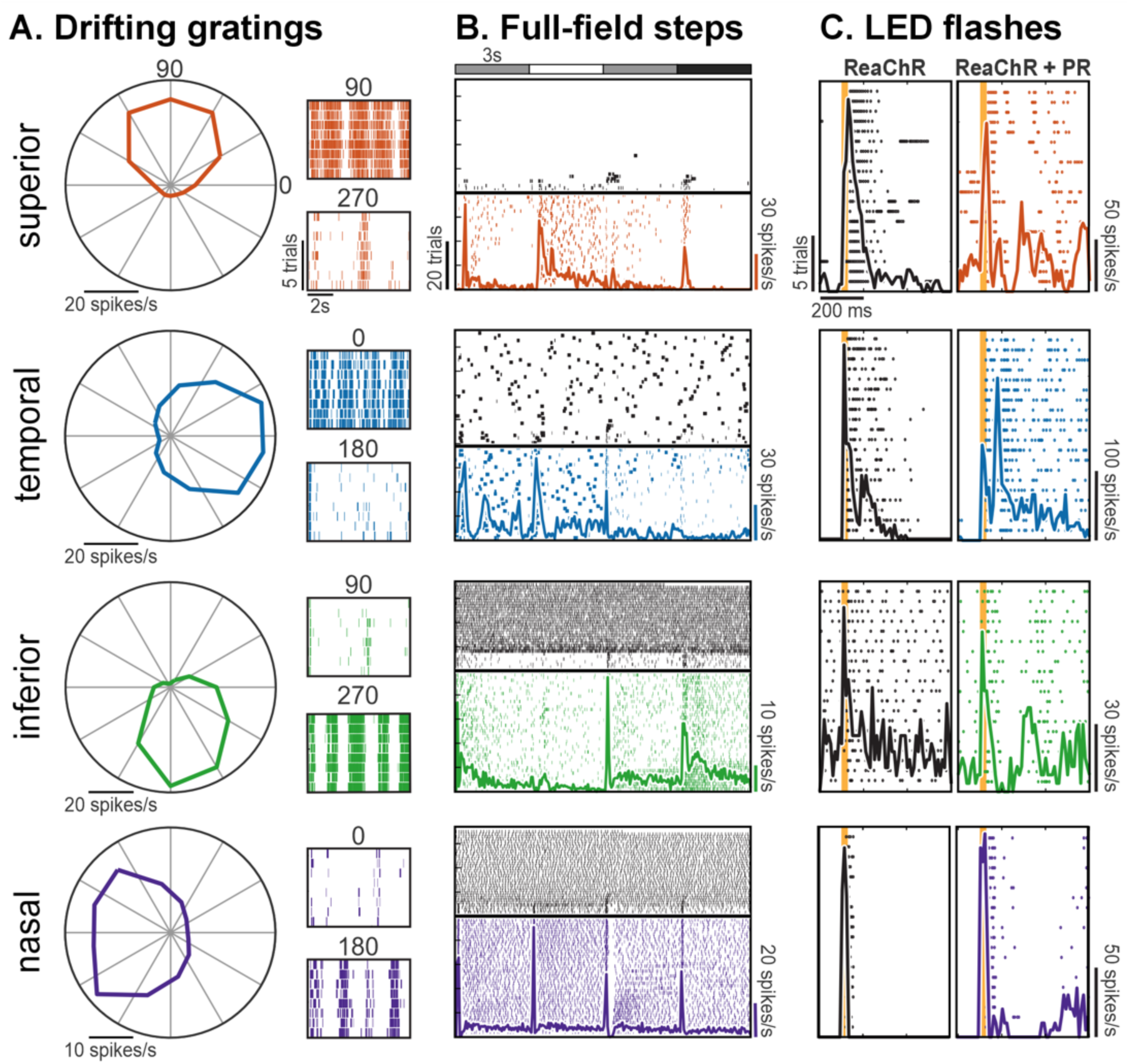
Direction-selective RGCs in rat project to SC. (**A**) Responses to drifting gratings for four direction-selective RGCs sensitive to each of the four cardinal directions. Gratings were drifted in 12 directions to generate the polar plots, the response magnitudes for motion in each direction. Spike rasters show the responses to the indicated preferred and null directions. (**B**) Spike rasters from full-field light steps that transitioned from gray-to-white-to-gray-to-black every 3 seconds (top). The colored portion of the rasters indicates trials prior to wash-in of drugs to block photoreceptor-mediated responses (see Methods), which were applied at the time indicated by the horizontal black line. Colored curves are peri-stimulus time histograms calculated during this period. Shortly after drug wash-in, spiking activity became uncorrelated with the full-field flashes, though spontaneous spiking varied across cells. (**C**) *Left column:* responses to a 100 ms 565 nm LED flash (orange bar) in the presence of drugs to block photoreceptor-mediated responses. Resulting spikes are mediated by ReaChR expression. *Right column*: responses to 565 nm LED flashes following drug wash-out, thus driven both by ReaChR and photoreceptors (PR). Flash strength was ∼10^14^ photons/mm^2^ to drive ReaChR.

**Supplemental Figure 3.**
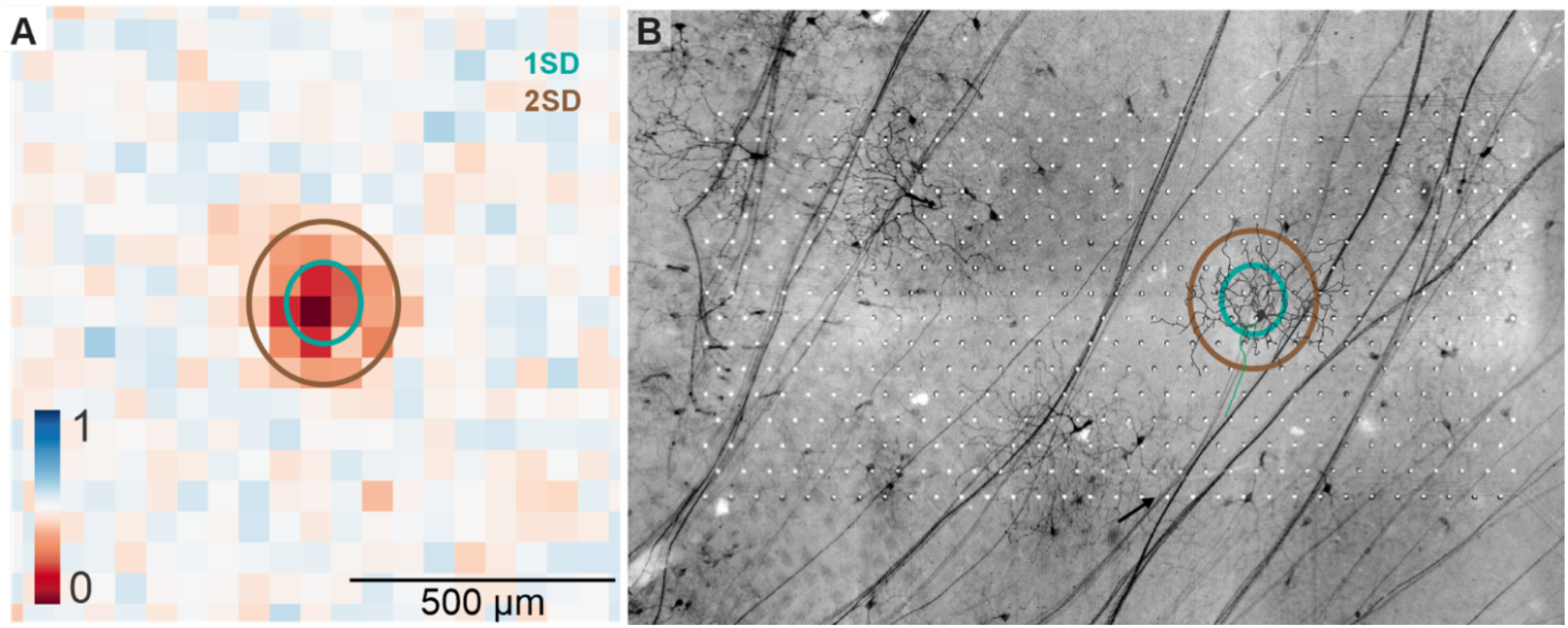
Spatial receptive field compared to morphology of example ReaChR-positive RGC. **(A)** One and two standard deviation (1-SD, teal; 2-SD, brown) contours of the RF center obtained from a difference-of-Gaussians fit are overlaid onto the SRF heatmap obtained from the STA to checkerboard noise. **(B)** The same Gaussian fit contours are overlaid onto a photomicrograph of the retina on the MEA. The traced neuron from Fig. 7H is overlaid.

**Supplemental Figure 4.**
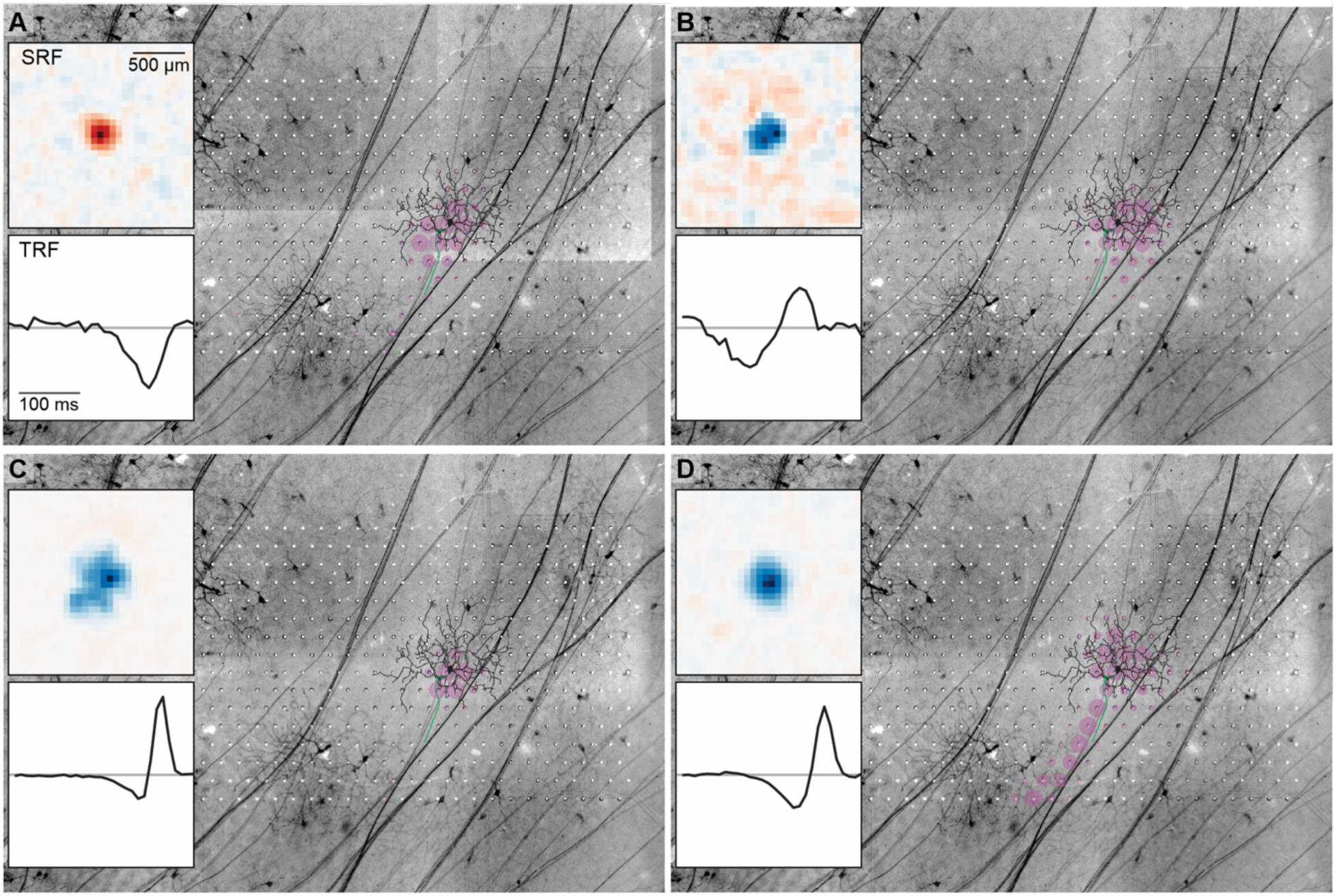
Example selection process when multiple EIs overlap a labeled RGC. **(A)** EI (pink) of the ReaChR-positive cell superimposed onto a photomicrograph of the live retina on the MEA. The tracing of the cell (Fig. 7H) is overlaid on top. Inset: spatial receptive field (SRF) and temporal receptive field (TRF). **(B-D)** Same as A, but for three RGCs with the next-best EI matches to the RGC morphology. Although the EIs for cells in B-D heavily overlap the labeled RGC, each exhibits an ON-polarity RF, which does not match the OFF-stratifying dendrites of the labeled RGC. Furthermore, none of the corresponding cells for EIs in B-D exhibited ReaChR-mediated responses to the 565 nm LED flashes. All EIs shown are calculated from spikes recorded during the checkerboard noise stimulus.

**Supplemental Figure 5.**
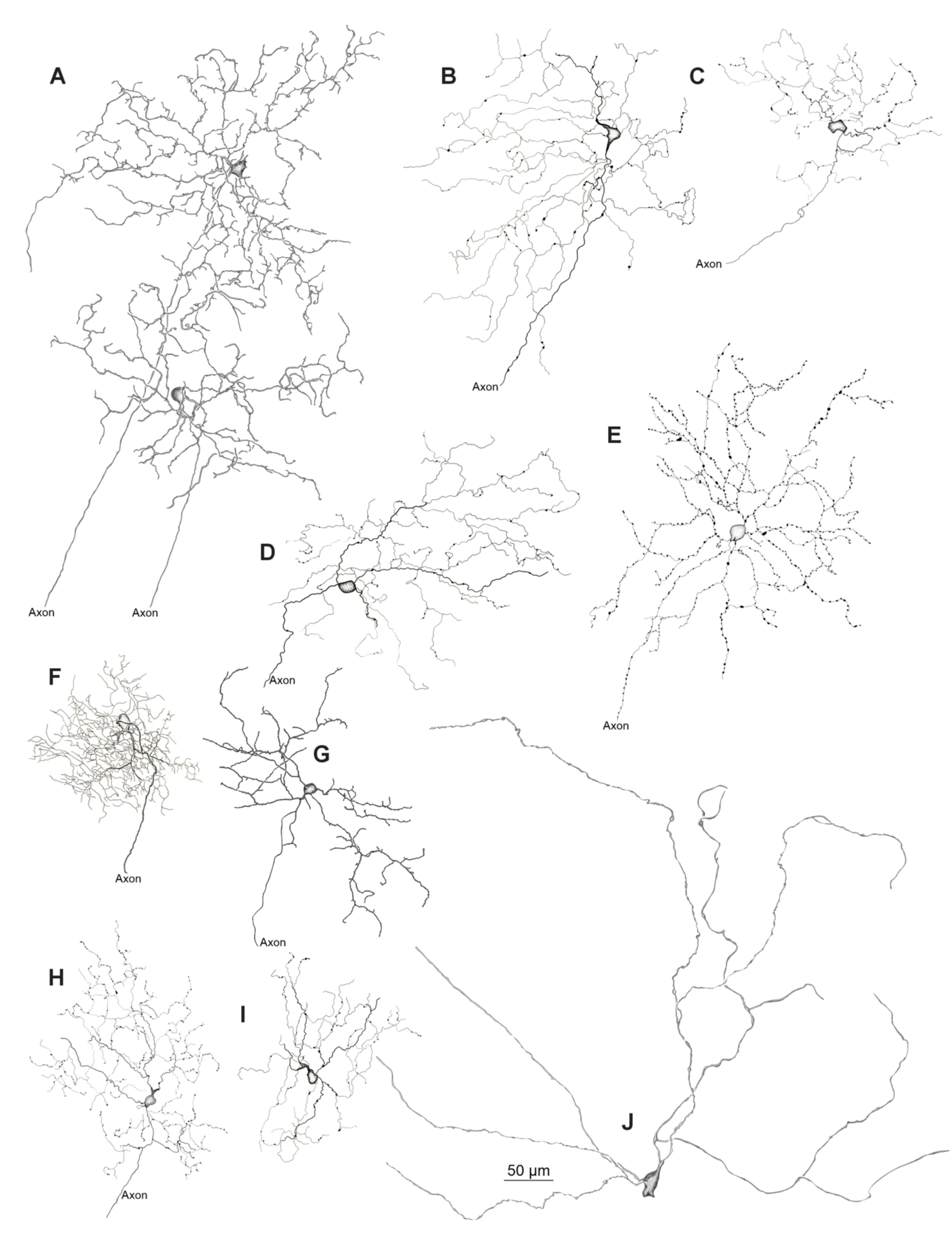
Morphological reconstructions of AAV2retro-transduced retinotectal RGCs. Each immunofluorescently amplified RGC (A, F, G) was drawn from 60x confocal scans (30-90 z-stacks) using the FIJI plugin Neuroanatomy. RGCs amplified using immunohistochemistry (B, C, D, E, H, I, J) were hand-drawn.

## ACKNOWLEDGEMENTS

We thank Jessi Cruger and Peter Saloupis for their assistance with fundoscopy. This work was supported by generous funding through a Duke Institute for Brain Sciences (DIBS) Incubator Award (G.D.F.; M.A.S.), NIH R01 EY034004 (G.D.F.; M.A.S.), K99 EY032119 (S.R.), NIH P30 EY005722 Core Grant for Vision Research at Duke University, and the NIH P30 EY000331 for Vision Research and an Unrestricted Grant from Research to Prevent Blindness at UCLA.

## AUTHOR CONTRIBUTIONS

**M.O.B.:** Conceptualization; Data curation; Formal analysis; Funding acquisition; Investigation; Methodology; Project administration; Supervision; Validation; Visualization; Writing—Original draft; Writing—review & editing**; A.M.R.:** Data curation; Formal analysis; Investigation; Methodology; Project administration; Software; Supervision; Validation; Visualization; Writing – original draft; Writing – review & editing; **T.B.D.:** Formal analysis; Investigation; Methodology; Writing – review & editing; **G.M.K.:** Formal analysis; Investigation; Methodology; Writing—Original draft; **E.S.:** Data curation; Formal analysis; Methodology; Software; Resources; Validation; Visualization; Writing – original draft; Writing – review & editing; **C.H.:** Formal analysis; Investigation; Methodology; **D.R.R.:** Visualization; **A.G.:** Visualization **S.R.:** Conceptualization; Funding acquisition; Investigation; Methodology; Software; Validation; Writing – original draft; Writing – review & editing; **K.R.:** Resources; **M.A.S.:** Funding acquisition; Project administration; Supervision; Writing – review & editing; **G.D.F.:** Conceptualization; Funding acquisition; Investigation; Methodology; Project administration; Resources; Supervision; Writing – review & editing.

## ABBREVIATIONS

AAV: adeno-associated virus
cMRF: central mesencephalic reticular formation
CRF: contrast response functions
DpG: deep gray layer of the superior colliculus
DpWh: deep white layer of the superior colliculus
EI: electrical image
GCL: ganglion cell layer
IC: inferior colliculus
INL: inner nuclear layer
InG: intermediate gray layer of the superior colliculus
IPL: Inner Plexiform Layer
InWh: intermediate white layer of the superior colliculus
ISI: inter-spike interval distribution
MEA: multi-electrode array
MGN: medial geniculate nucleus of the thalamus
OT: nucleus of the optic tract
Op: optic nerve layer of the superior colliculus
OSI: orientation-selective index
PAG: periaqueductal gray
PC: posterior commissure
POD: post-operative day
PT: pretectum
ReaChR: red-shifted channelrhodopsin
RF: receptive field
RGC: retinal ganglion cell
SRF: spatial receptive field
SuG: superficial gray layer of the superior colliculus
TRF: temporal receptive field
Zo: zonal layer of the superior colliculus

## STAR METHODS

### Viral Vectors

AAV2retro-CAG-ReaChR-mCitrine was produced at UNC neuroscience center/brain initiative neurotools vector core. Titers ranged from 10^12^ to 10^13^ GC/ml.

### Animals

Long-Evans rats (Charles River Laboratories), aged 6–9 weeks and of both sexes, were used for the experiments. All procedures adhered to NIH guidelines and were approved by the Duke University Institutional Animal Care and Use Committee.

### SC Injections

Rats were positioned sternally in a stereotaxic apparatus (Model 940, David Kopf Instruments, Tujunga, CA). Anesthesia was maintained with 1-3% isoflurane mixed with oxygen. Carprofen (3 mg/kg; SQ) was administered as a pre-operative analgesic. Bupivacaine (∼0.5 ml; SQ) was subcutaneously applied before the first incision and after the final suture.

A midline incision was made, and the skull was bilaterally trephined at -6.3 mm (anterior-posterior) and ±1.5 mm (medial-lateral) relative to Bregma. Injections were performed using a 10 µL Neuros Hamilton syringe (Model 1701 RN, 33 gauge, Point Style 3, Hamilton Company, Reno, NV; Part/REF #65460-05) mounted to a microsyringe pump (UMP3T-1, World Precision Instruments, Sarasota, FL). A single penetration was made in each hemisphere, and the needle tip was advanced to a depth of -3.6 mm from the cortical surface. A 333 nL volume of viral solution was dispensed at a rate of 200 nL/minute. The needle remained in place for 5 minutes to allow for viral diffusion. This process was repeated at two additional depths of -3.3 mm and -3.1 mm. After the final injection, the needle was left in place for 10 minutes to facilitate further viral diffusion before being fully retracted and moved to the contralateral hemisphere to repeat the injection procedure. In some cases, single injections of 333 nL were made within specific layers of the SC. In this case, a single injection was made followed by a ten-minute wait period. Once the injections were completed, the skin was reapproximated and sutured. For post-operative care, a second dose of Carprofen (3 mg/kg; SQ) was administered the following day.

### Fundus imaging

To assess the location and timeline of retrograde transgene expression in the retina, fluorescence fundus ocular imaging was conducted on rats following viral injections into the SC. The rats were initially anesthetized with ketamine (40 mg/kg; IP) followed by dexdomitor (0.5 mg/kg; IP). Once anesthetized, 1% atropine sulfate and 0.5% proparacaine hydrochloride eye drops were applied to both eyes to dilate the pupils and reduce corneal sensitivity. The animal was then positioned to align the eye with the imaging objective of a fundus microscope (Heidelberg SPECTRALIS HRA + OCT, Heidelberg Engineering, Heidelberg, Germany). The blood vessels and optic nerve were visualized by using the infrared-reflectance single imaging mode and then the focus imaging plane was set to near the surface of the retina. Next, the imaging mode was switched to fluorescence (excitation wavelength 488 nm). Digital photomicrographs were taken using the Fluorescein Angiography single imaging mode, with the Automatic Real Time feature enabled and normalization turned off. The retina was imaged region by region, refocusing as necessary to capture the entire retina. After completing the imaging of both eyes, anesthesia was reversed using antisedan (atipamezole, 0.5 mg/kg; IP), and the animal was returned to its home cage.

### Multielectrode array (MEA) recording and spike sorting

The rats were anesthetized in an induction box using a 5% isoflurane-oxygen mixture. After the animal was immobile, it was transferred to the stereotaxic instrument and injected with Ketamine/Xylazine (150 mg/kg/10 mg/kg; IP). Once completely areflexic, the eyes were enucleated under ambient room light. The eyes were immediately placed in an oxygenated solution of Ames’ media and transferred to a dark room. Both eyes were then hemisected and vitreous removed under infrared illumination using infrared goggles. To dark adapt the retina, the posterior half of the eye with retina attached to the pigment epithelium was placed in an incubation dish containing oxygenated Ames’ solution at 32°C for 45 - 60 minutes. Next, a region of the retina that was found to contain fluorescently labeled RGCs from fundus imaging was identified, and a ∼3 mm x 1.5 mm biopsy was collected. This piece of retina was placed RGC side down on the MEA (512 electrodes spanning an area of 0.9 x 1.8 mm (Litke et al., 2004), 60 µm pitch). Extracellular voltages were sampled at 20 kHz, filtered, and stored for spike sorting.

Spike sorting was achieved using at least two independent methods: (1) a custom-written spike sorter called Vision (Shlens et al., 2006) and (2) YASS (Lee et al., 2020) or (3) Kilosort (2.5 and 4)(Pachitariu et al., 2016; Pachitariu et al., 2023). Vision utilized principal component analysis and a mixture of Gaussian models for spike clustering. YASS and Kilosort methods are described in detail in previous work (Pachitariu et al., 2016; Lee et al., 2020; Pachitariu et al., 2023). RGCs generating few spikes (<100 spikes) and those with high contamination (>10% in 1.5 ms spike refractory period) were discarded. RGCs were retained for further analysis if they exhibited stable responses across multiple stimulus conditions and satisfied cross-validation of electrophysiological images.

### Visual stimulation

Visual stimuli were created in MATLAB (Mathworks, Natick, MA) using an OpenGL framework and delivered using a gamma-corrected micro-OLED (Emagin SVGA+XL, Santa Clara, CA). The stimulus was focused onto the photoreceptor outer segments using a 100 mm focal length bi-convex lens (Thorlabs, LB1630) and an objective (Nikon, 4x Plan Fluor, MRH00045) mounted on an inverted microscope (Nikon, Eclipse Ti-2). The photon flux was measured using a photometer (UDT Instruments, Gamma Scientific) and was converted to photoisomerization rate per photoreceptor (R*/photoreceptor/s)(Baylor et al., 1984; Field and Rieke, 2002; Yu et al., 2017). The following stimuli were used: (1) checkerboard binary white noise (check size 60 mm x 60 mm) refreshing every 1 or 2 frames, with frame rate 60.35 Hz, (2) gratings drifting along 12 directions with spatial frequency 8.4*10^-4^ m^-1^ and temporal frequencies 1 Hz and 4 Hz (corresponding to speeds 2.8 cyc/mm and 11.2 cyc/mm respectively) at 95% Weber Contrast, and (3) full-field light steps (0%, 50%, 100% contrasts) with each step lasting 3 seconds.

### Receptive field estimation

Spatiotemporal receptive fields (RFs) were estimated using previously described methods (Gauthier et al., 2009; Roy et al., 2021). Briefly, the spike-triggered average (STA) was calculated using 500 ms of stimulus frames preceding each spike. Stimulus pixels with values exceeding 3.5 standard deviations of all stimulus pixels were selected as putative pixels within the RF. The temporal evolution of stimulus pixel values at these locations were averaged to estimate the temporal RF. To estimate the spatial RF, the inner product of the STA and the temporal RF was computed, which collapsed the STA across time. The spatial RF was fit with a spatial difference of Gaussians and the temporal RF was fit with a cascade of temporal filters (Chichilnisky and Kalmar, 2002; Ravi et al., 2018). The spatial RF size was quantified using the geometric mean of the standard deviations along the major and minor axes of the Gaussian fit. The phasic index (PI) was calculated from the temporal RF fit as follows:

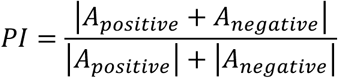

Where A_positive_ and A_negative_ are the area under the positive temporal kernel and the area above the negative temporal kernel, respectively. For visualization, the spatial RF was filtered with a circular Gaussian filter (SD = 0.75 stimulus pixels),

### Direction- and orientation-selective responses

Responses to gratings drifting along 12 directions (30 degrees apart) were used to calculate a direction-selective index (DSI) and an orientation-selective index (OSI). The total number of spikes over N trials for each direction of movement was used to calculate the DSI and the OSI:

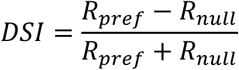

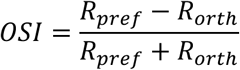

Where R_pref_ is the response in stimulus direction closest to the preferred direction, R_null_ is the response in the opposite stimulus direction, and R_orth_ is the response in the orthogonal stimulus direction. RGCs with DSI greater than 0.3 were selected as direction-selective. RGCs with OSI greater than 0.3 and DSI less than 0.3 were selected as orientation-selective.

### Optogenetic stimulation

RGCs that express ReaChR can be activated by both visual stimulation of photoreceptors and activation of ReaChR using 565 nm light (Lin et al., 2013). To isolate spikes generated solely from ReaChR activation, photoreceptor-driven input to the ON and OFF bipolar cells was pharmacologically blocked using a cocktail of L-AP4 (mGluR6 agonist, 100 µM; Tocris Bioscience, Cat. # 0103), DL-AP5 (NMDA antagonist, 100 µM; Tocris Bioscience, Cat. # 0105), and CNQX (AMPA and kainate antagonist, 100 µM; Tocris Bioscience, Cat. # 0190)(Slaughter and Miller, 1981; DeVries, 2000; Wong, 2012; Puthussery et al., 2014). Spiking responses of RGCs to full-field light steps were first measured under control condition (Ames’ solution) for at least 300 s, and then the drug cocktail was washed in. The responses to light steps were abolished in 3-5 minutes after drug wash-in (**Fig. 4D**). Next, a 565 nm LED (Thorlabs, M565L3) was used to generate sequences of brief flashes of light to photoactivate ReaChR-expressing RGCs. Flash durations of 10, 30, 100, and 200 ms at intensities of 0.25, 0.74, 2.75, 4.64 x 10^8^ R*/rod/s were each delivered 20-30 times, with at least a one-minute interval between each flash strength. The flash intensities were higher than the intensity threshold for ReaChR responses in the retina: ∼ 0.2 x 10^8^ R*/rod/s (Sengupta et al., 2016). Finally, the drug cocktail was washed out with Ames’ solution for 10 minutes, and then response measurements to the LED flash sequences were repeated.

### ReaChR cell identification

ReaChR-expressing RGCs were identified based on their responses to 565 nm LED flashes delivered at nine flash strengths generated by combining the intensities at different flash durations. For each flash strength, the trial-averaged ReaChR signal (30 trials) was computed as the mean spike rate within 100 ms of LED onset from which the baseline spike rate measured 2–3 seconds after stimulus onset was subtracted (**Supp.** Fig. 1A). These nine values formed the intensity-response function for each cell (**Supp.** Fig. 1B, C). These functions were used as vectors in a principal component analysis (**Supp.** Fig. 1D). The first two principal components (PCs) accounted for 92.2% (PC1) and 3.7% (PC2) of the variance. With PC1 dominating the variance, intensity-response functions were projected along this PC and binned into a histogram. We labeled these projection values the Phototagged Response Index (PRI). These histograms were approximately Gaussian with a long tail of positive values in ReaChR-expressing retinas; ReaChR-negative retinas were approximately Gaussian with no tails. Thus, these histograms were fit with a Gaussian distribution and ReaChR-positive RGCs were classified as cells exceeding a threshold of 4 standard deviations above the mean of this distribution. This threshold choice resulted in no false-postive ReaChR-expressing cells in retinas without ReaChR expression.

### Electrical images (EIs)

For each neuron, the EI was computed as the STA of the electrical activity across the MEA, serving as a spatial "footprint" to track cells across stimulus and drug conditions (Litke et al., 2004; Petrusca et al., 2007; Ruda et al., 2020). To construct the EI, we averaged analog waveforms from all 512 electrodes time-locked to the spikes of a neuron. Waveform polarity and shape distinguished cellular compartments: somatic (biphasic: downward-upward), dendritic (biphasic: upward-downward), and axonal (triphasic) signals (Petrusca et al., 2007; Wu et al., 2023). EIs were aligned to epifluorescence images to confirm ReaChR-positive cell identity.

### Tracking RGCs across stimulus conditions

The EI was estimated for each RGC, separately for responses to visual stimuli and stimulation from 565 nm LED. The spatial EIs from different stimulus conditions were correlated and a threshold correlation of 0.9 was used to confirm that the EIs originated from the same RGC (**Fig. 6; EIs**)(Field et al., 2009; Yao et al., 2018).

### Histology

#### Retina immunofluorescence

The retinas were fixed in 4% paraformaldehyde (PFA) for 45 minutes at room temperature and then stored in phosphate buffered saline (PBS) + 0.02% sodium azide (Santa Cruz Biotechnology, Cat # sc-296028) at 4°C. For whole-mount staining, the retinas were incubated in 5% normal donkey serum (Jackson Immunoresearch, Cat # 017-000-121) in 0.75% Triton X-100 (VWR Life Sciences, Cat # 97062-208) and 0.75% Tween 20 (Sigma-Aldrich, Cat # P1379) overnight at 4°C, then in primary antibodies for 5 days at 4°C, followed by secondary antibodies overnight at 4°C. The retinas were washed in PBS + 0.02% sodium azide three times over 3 hours at RT, then incubated in Hoechst 33258 (1:1000, Thermo Scientific, 62249) for 15 minutes. Retinas were flat mounted on glass slides with coverslips and stored at 4°C for confocal imaging.

#### Antibody concentrations

Primary antibodies included chicken anti-GFP (1:1000; Invitrogen A10262) and goat anti-ChAT (1:500; EMD Millipore Cat # AB144P). Secondary antibodies were conjugated to Alexa Fluor 488 (1:500; Invitrogen, A78948) and Alexa Fluor 555 (1:500; Invitrogen, A32816).

#### Retinal immunohistochemistry

After fixation (described above), endogenous peroxidase activity was blocked by incubating retinas in 0.3% hydrogen peroxide PBS solution for 30 minutes. The retinas were transferred to a blocking solution containing 1.0M PBS, 0.75% Triton-X 100, 0.75% Tween-20, and 5% normal donkey serum and incubated overnight at 4°C. Following this, chicken anti-GFP IgY (1:500) was added to the blocking solution, and the retinas were again incubated overnight. The retinas were rinsed, then placed in a solution containing 5% normal donkey serum and biotinylated donkey anti-chicken IgY (1:500, Invitrogen; A16003) as the secondary antibody and incubated overnight. After rinsing with PBS, the retinas were incubated in a 1% Avidin-Biotin horseradish peroxidase solution (Vector Labs; PK-6100) for 1 hour. They were then rinsed again and transferred to a solution containing 1% cobalt chloride, 1% nickel ammonium sulfate, and diaminobenzidine tetrahydrochloride for 25 minutes. To catalyze the chromogen reaction, 3% hydrogen peroxide was added to this solution, and the retinas were incubated for an additional 45 minutes. Finally, the retinas were rinsed, flat-mounted on glass slides, and coverslipped.

#### RGC Morphology

Retinas were examined using a Nikon Eclipse Ti2 confocal microscope, controlled by NIS-Elements AR software (Nikon Instruments Inc., Melville, NY, U.S.A.). A panoramic image was first captured to encompass the entire dendritic morphology. Following this, a Z-stack was obtained at 0.175-micron intervals, covering all viral-mediated labeling observed along the Z-axis. For manual reconstruction of RGCs, the confocal Z-stacks were imported into CorelDRAW, where each neuronal feature was traced as vectored lines on separate layers. The line thickness was adjusted to match the average diameter of the traced dendrites, while the axon was given a heavier line weight to make it stand out. For semiautomated reconstructions, confocal Z-stacks were imported into the FIJI plugin, SNT Neuroanatomy (Arshadi et al., 2021), for more efficient tracing of neuronal structures.

#### SC immunofluorescence

To confirm injection locations, the brains were extracted for histological confirmation. Immediately following enucleation, each animal was exsanguinated by transcardial perfusion. The vasculature system was first flushed with PBS, followed by a 4% PFA wash. The brain was dissected and postfixed in fresh 4% PFA for ∼24 hours. Next, PFA was rinsed out, and the brain was moved to a 30% sucrose in PBS solution for cryoprotection. The brain was sectioned at 75 µm on a freezing stage sliding microtome (American Optical).

After sectioning, the brain was divided into three series. The first series was reserved for epifluorescence imaging. Free floating sections were counterstained with Hoechst 33342 (1:1000; ThermoFisher, 62249) in 7.4 pH 0.1M PBS for 15 minutes and then washed with PBS before being mounted. Tissues forming the other two series were immunofluorescently amplified. Tissue was permeabilized using 0.5% Triton-X 100 solution, blocked with 1% BSA, then incubated for ∼12 hr in goat anti-GFP (1:800; Rockland, 600-101-215). Next, the tissue was rinsed in PBS and placed into a PBS solution containing donkey anti-goat Alexa Fluor 488 (1:200; ThermoFisher, A-11055) and 10% normal donkey serum (Jackson ImmunoResearch, 017-000-121) for 3 hours at room temperature. The tissue was then counterstained with Hoechst 33342. In all series, sections were mounted on 10% gelatinized slides, dehydrated in graded concentrations of ethanol, then xylene, and coverslipped with Cytoseal 60 (Fisher Scientific, 23244257)(Daw et al., 2023).

### Key Resources Table

| REAGENT OR RESOURCE | SOURCE | IDENTIFIER |
| --- | --- | --- |
| Hoechst 33342 | ThermoFisher | 62249 |
| Hoechst 33258 | ThermoFisher | H1398 |
| Avidin & Biotin | Vector Labs | PK-6100 |
| **Bacterial and virus constructs** |  |  |
| AAV2retro-CAG-ReaChR-mCitrine | UNC-Neurotools |  |
| **Antibodies** |  |  |
| Goat Anti-GFP | Rockland | 600-101-215 |
| Chicken Anti-GFP | Invitrogen | A10262 |
| Goat Anti-ChAT | EMD Millipore | AB144P |
| Donkey Anti-Goat AF 488 | ThermoFisher | A-11055 |
| Donkey Anti-Chicken AF 488 | Invitrogen | A78948 |
| Donkey Anti-Goat AF 555 | Invitrogen | A32816 |
| Donkey Anti-Chicken IgY Biotinylated | Invitrogen | A16003 |
| **Softwares and algorithms** |  |  |
| FIJI | NIH | https://imagej.net/software/fiji/ |
| MATLAB | Mathworks | https://www.mathworks.com |
| CorelDRAW | Alludo | https://www.coreldraw.com/en/ |
| **Experimental models (organisms & strains)** |  |  |
| Rat (Long-Evans) | Charles River Laboratories | 006 |

